# Locus Coeruleus firing patterns selectively modulate brain activity and dynamics

**DOI:** 10.1101/2022.08.29.505672

**Authors:** Christina Grimm, Sian N. Duss, Mattia Privitera, Brandon R. Munn, Stefan Frässle, Maria Chernysheva, Tommaso Patriarchi, Daniel Razansky, Nicole Wenderoth, James M. Shine, Johannes Bohacek, Valerio Zerbi

## Abstract

Noradrenaline (NA) release from the brainstem nucleus locus coeruleus (LC) changes activity and connectivity in neuronal networks across the brain, thus modulating multiple behavioural states. NA release is mediated by both tonic and burst-like neuronal LC activity. However, it remains unknown whether the functional changes in downstream projection areas depend on these firing patterns. Using optogenetics, pupillometry, photometry, and functional MRI in mice, we show that tonic and burst LC firing patterns elicit brain responses that are specific to the activation frequency and temporal pattern. Tonic activation of the LC evokes nonlinear responses in prefrontal, limbic, and cerebellar regions, in line with the proposed inverted-U relationship between LC activity and behaviour. We further demonstrate that LC activity enhances network integration and acts as a facilitator of brain state transitions, hence increasing brain flexibility. Together, these findings reveal how the LC-NA system achieves a nuanced regulation of global circuit operations.

## Introduction

The brain’s capacity to rapidly shift between different modes of information processing while working with limited resources supports an array of behavioural goals and contextual demands in a dynamic environment. The brain is thus required to orchestrate changes in the state of its neural network to manage the efficiency of specific circuit interactions. To make the process deterministic and not random, it has been theorised that the brain must spend a certain amount of energy to engage in a state transition, the level of which can be modulated internally, much like in a catalysed system ^1-3^. Such a prerequisite raises a fundamental question: which neural structures and mechanisms can modulate the probability of state shifts, while stably balancing ongoing behaviour and rapid adaptation? Over the past years, several theories have pointed to the locus coeruleus - noradrenaline (LC-NA) system and its neuromodulatory actions as a potential candidate ^4-6^. Located in the pontine brainstem with wide projections throughout the whole brain, the LC is ideally situated to accommodate the critical demands of an ever-changing environment ^1, 7^. It is the principal site of synthesis of the neuromodulator NA and recognised to contribute to many central nervous system functions, including sensory processing, modulation of arousal, nociception, sleep/wake cycles, cognition and stress responses ^8^.

A commonly held view is that the inherent patterns of LC activity are key drivers of various behavioural processes that can operate on sub-second timescales (e.g., orienting attention) up to minutes or hours (e.g., vigilance, stress) ^9^. Although mostly referred to as distinct modes of firing, tonic and phasic firing likely present extremes of a continuum of LC function ^4^. In monkeys, rats and mice, spontaneous discharges across a range of relatively low frequency (0.5-8 Hz; tonic activity) can be coarsely related to arousal levels ^7, 10-13^. In contrast, brief bursts of activity (2-4 spikes at 10-25 Hz) occur in association with salient or novel stimuli and decisions ^12-15^ and are thought to facilitate behavioural responses by increasing the neural gain in target regions at task-relevant timepoints ^4, 16^. Analyses and modelling of functional MRI (fMRI) signals during pharmacological challenges ^17, 18^, behavioural tasks ^19, 20^ or even at rest ^21-25^ are in agreement with this view and have provided further evidence that LC activity levels influence the dynamics of brain states ^26^. For example, periods of increased arousal, compatible with burst firing of the LC, rapidly recruit the prefrontal and posterior parietal cortex and result in an improvement of executive function in humans ^23^. Instead, exposing subjects to strong stressors (compatible with high-tonic LC activity) causes rapid disengagement of prefrontal areas and induces the mobilisation of the amygdala and sensory-motor cortices to promote hypervigilance and threat detection, albeit at the expense of executive control ^23, 27-29^. More recent analyses from high-resolution (7T) resting-state human fMRI datasets further established a correlation between LC activity and the likelihood of a network’s state transition ^30, 31^.

The involvement of NA-signalling in the control of the dynamics of connectivity states has been confirmed using the pharmacological blockade of adrenergic receptors ^28^. However, the causal contribution of the LC to shaping brain dynamics has only been addressed recently, as chemogenetic modulation of the LC in mice was found to be sufficient to trigger a rapid remodelling of network connectivity ^32, 33^. This work also revealed that adrenergic receptor distribution likely explains some of the regional specificity that noradrenergic neuromodulation can achieve, despite the diffuse projection pattern of the LC ^32^. However, due to technical constraints of chemogenetics, these studies were not able to address the key question if (and how) different activity patterns of the LC can shape brain network dynamics.

In this work, we set out to gain a fundamental understanding of the neural bases of LC-guided mechanisms in brain activity changes and dynamic network reconfigurations. We used optogenetics to mimic physiologically relevant excitatory patterns of tonic and burst LC firing while performing pupillometry, measuring local NA release with fibre photometry or recording whole-brain fMRI responses. Our data show that LC-NA output changes whole-brain activity in a manner that is dependent on both firing pattern and frequency. Using a low-dimensional landscape framework, we further demonstrate topographical changes in cortical and thalamic energy landscapes that support the proposal that the LC-NA system acts as a catalyst for brain state transitions. Furthermore, generative modelling of effective (directed) connectivity revealed frequency-dependent alterations in functional coupling amongst key target regions of the LC: the hippocampus, the amygdala, and the prefrontal cortex. Taken together, these results present a causal view of the LC-NA system in shaping cortical and sub-cortical dynamics through its distinct firing patterns.

## Results

### Whole-brain activity changes with levels of tonic LC firing

LC neurons have been shown to spontaneously discharge across a continuum range of relatively slow rates (0.01-5 Hz; tonic activity) coarsely related to levels of arousal ^10, 11^. However, how these different tonic LC firing frequencies affect whole-brain activity is yet to be understood. To achieve spatially and temporally precise modulation of LC noradrenergic neurons, we unilaterally transfected the right LC of DBH-iCre mice (n(male)=22; n(female)=7) with an AAV construct carrying the optogenetic actuator channelrhodopsin-2 (ChR2) and implanted an optical cannula above the target site. We assessed the physiological impact of optogenetically evoked tonic LC firing using pupillometry ^32, 34^. Briefly, a 2-minute baseline recording was followed by different laser stimulations as shown in **Figure 1A**. We chose 3Hz, because several studies had suggested that this intermediate level of LC activity is associated with optimal task performance ^4, 35-37^. We chose 5 Hz, because previous work had shown that tonic 5Hz stimulation phenocopies the effects of strong acute stressors ^38^, and that higher stimulation frequencies (10Hz) might actually lead to LC-exhaustion ^26^ or LC inhibition ^39^. Further, NA release scales linearly with LC activity up to around 6Hz ^24^. We first confirmed that unilateral optogenetic stimulation of LC noradrenergic neurons resulted in pupil dilation at both tonic LC stimulation frequencies compared to sham stimulation (**Figure 1B)**. As previously described ^34^, we found that activating the LC with 5 Hz stimulation, changes in pupil size were significantly greater than at 3 Hz (mean±SD(3 Hz) = 14.24±6.1, mean±SD(5 Hz) = 16.47±7.3, paired t-test; t(23)=3.7, p=0.0012) (**Figure 1C**), suggesting that increasing intensities of tonic LC-NA activity trigger a graded pupillary response. Differential release of NA was further validated using fibre photometry recordings in the hippocampus (HP), a brain region receiving noradrenergic input exclusively from the LC ^40^. We found that the change in fluorescence of the NA-sensor GRAB_NE1m_ ^41^ was greater in response to tonic 5 Hz stimulation compared to tonic 3 Hz LC stimulation (mean±SD(3 Hz) = 2.98±1.88, mean±SD(5 Hz) = 4.95±3.23, paired t-test; t(5)=3.26, p=0.022).

**Figure 1.**
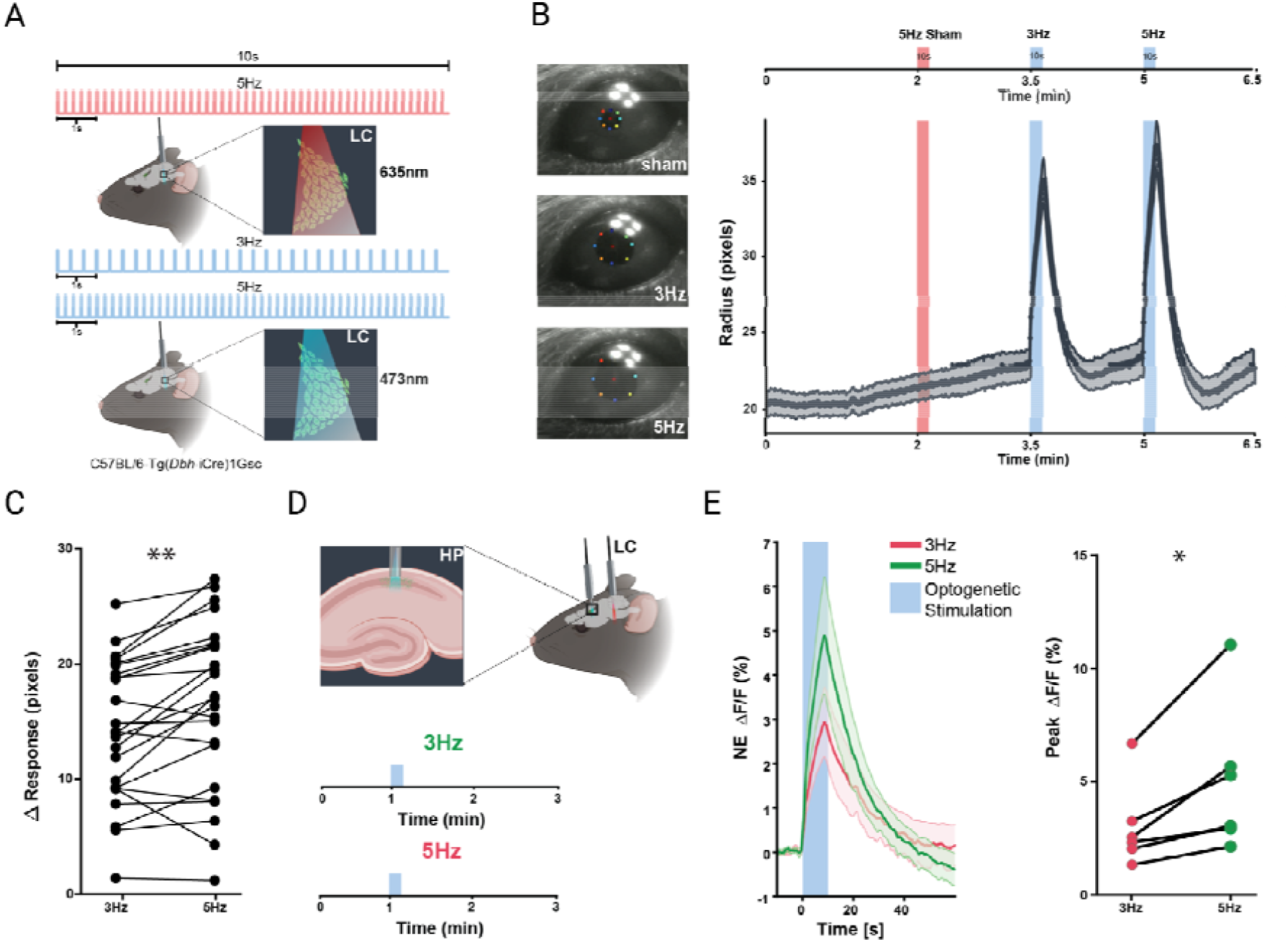
Physiological effects of tonic LC activation. **A** Schematic of 3 Hz and 5 Hz tonic, blue laser-light stimulation and red laser-light, sham stimulation protocols for pupillometry. The protocol included a 2-minute baseline recording followed by 10 second stimulations: 1) 5 Hz at 635nm as control, 2) 3 Hz at 473nm, and 3) 5 Hz at 473nm. **B** Left: representative pictures during sham, 3Hz, and 5Hz LC stimulation. Right: pupil traces at sham, 3 Hz tonic, and 5 Hz tonic LC stimulation (n=24, mean±SEM). **C** Statistical comparison of the pupil response to 3 Hz and 5 Hz tonic LC stimulation. 5 Hz LC stimulation induces a significantly greater pupil dilation than 3 Hz (mean±SD(3 Hz) = 14.24±6.1, mean±SD(5 Hz) = 16.47±7.3, paired t-test; t(23)=3.7, p=0.0012). **D** Schematic of optogenetic LC stimulation combined with fibre photometry in the HP. **E** Left: ΔF/F traces of GRAB_NE1m_ photometry recordings in response to tonic 3 Hz and 5 Hz LC stimulation (mean±SEM). Right: 5 Hz tonic LC stimulation triggers greater NA release compared to 3 Hz tonic LC stimulation (mean±SD(3 Hz) = 2.98±1.88, mean±SD (5 Hz) = 4.95±3.23, paired t-test; t(5)=3.26, p=0.022; n(male)=5, n(female)=1). HP, hippocampus; **p < 0.01; *p < 0.05

Next, we sought to assess whole-brain, time-locked effects of tonic LC firing by using a combined opto-fMRI approach with blue-light stimulation at 3 Hz and 5 Hz and a red-light control stimulation (**Figure 2A**). Pre-processed and normalised functional datasets were modelled with a General Linear Model (GLM), where the design matrix recapitulated the temporal parameters of the respective stimulation protocol to separate stimulus-induced signals from background activity. The experimental regressors in the design matrix were convolved with a custom hemodynamic response function (HRF) ^42^. This analysis revealed no traceable BOLD signal changes in sham stimulated mice (**Figure 2B**). For both 3 Hz and 5 Hz datasets, the BOLD signal in the targeted LC initially dropped, and then peaked towards the end of each 30 sec stimulation block (**Figure 2C-D**). As expected, no statistically significant brain-wide BOLD activation patterns emerged during sham stimulation (**Figure 2E, top panel**). In contrast, statistical activation maps of tonic stimulation at 3 Hz revealed increased BOLD signals in multiple regions directly innervated by the LC, including the infralimbic (IL) / prelimbic (PrL) and anterior cingulate area (ACC), thalamic nuclei including the mediodorsal nucleus (MD), the ventral posteriomedial nucleus (VPM) and the ventral posterolateral nucleus (VPL), the ventral hippocampus (HP) and the ipsilateral somatosensory barrel field cortex (SSCtx) (**Figure 2E, middle panel**). In agreement with our previous observation of increased network connectivity in the dorsal striatum upon chemogenetic LC activation, we detected robust BOLD signal increases in the bilateral dorsal striatum (CPu), despite its sparse innervation by LC axons ^32, 33^. Tonic LC stimulation at 5 Hz strongly increased BOLD signals in the ipsilateral somatosensory upper limb cortex and bilateral thalamic nuclei (**Figure 2E, bottom panel**). Instead, significant BOLD reductions were seen in the lateral septal nucleus (LS), the bilateral amygdala (Amy), the dorsal CPu, the periaqueductal gray area (PAG) as well as in the contralateral LC (**Figure 2E, top panel)**, where axonal projections from the optogenetically stimulated site can be found (**Figure S1**). Together, these data suggest that subtle differences in tonic LC firing frequency can evoke very distinct changes in brain-wide activity patterns.

**Figure 2.**
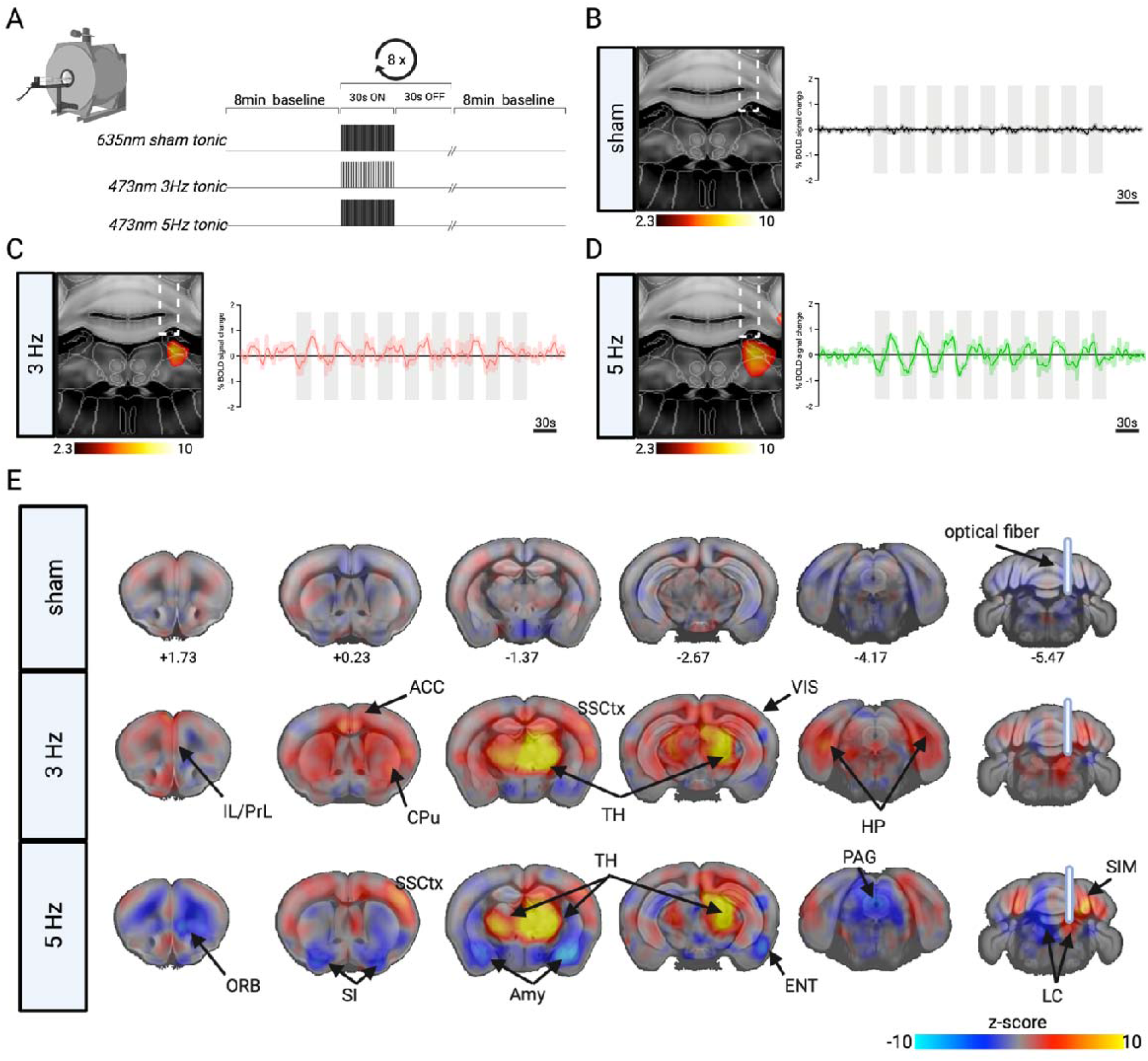
Local and brain-wide effects of tonic LC stimulation patterns. **A** Schematic of tonic, blue laser-light stimulation and red laser-light, sham stimulation protocols for combined opto-fMRI. **B-D** Mean time-series extracted from the optogenetically targeted, right LC of sham, 3Hz, and 5Hz datasets. The BOLD response evoked by LC stimulation shows an initial dip followed by a delayed increase most likely because NA release affects not only neural but also vasomotor activity. Time-series of sham tonic stimulation depicted in black, time-series of 3Hz tonic stimulation depicted in red, time-series of 5Hz tonic stimulation depicted in green. Grey bars represent 30s laser stimulation blocks. **E** Unthresholded general linear model (GLM) z-stat activation maps of tonic sham, 3 Hz tonic, and 5 Hz tonic stimulation of LC noradrenergic neurons. IL, infralimbic area; PrL, prelimbic area; ACC, anterior cingulate; Amy, amygdala; CPu, caudate putamen; ENT, entorhinal cortex; HP, hippocampus; LC, locus coeruleus; ORB, orbitofrontal area; PAG, periaqueductal gray; SI, substantia inominata; SIM, simple lobule; SSCtx, somatosensory cortex; TH, thalamus; VIS, visual area; VPM, ventral posteriomedial nucleus of thalamus. N(sham)=32; n(3Hz)=15; n(5Hz)=16.

### Different tonic LC firing rates evoke distinct activity and connectivity profiles amongst target brain regions

The GLM findings presented above (**Figure 2**) suggest that different firing patterns of the LC drive highly specific whole-brain activation patterns. This is in line with previous work showing that LC stimulation at 5 Hz induced strong place aversion, but no such effect was observed at 2 Hz stimulation ^38^. Similarly, the performance of monkeys on a visual discrimination task differed depending on small fluctuations in tonic firing frequency ^37^. It has thus been suggested that LC firing frequency follows the inverted-U (Yerkes-Dodson) relationship between arousal and task performance ^43^. To test whether we see evidence for such a relationship at the level of whole brain activation and to parse regional selectivity of LC firing across the tonic range, we directly compared BOLD activation maps of 3 Hz, 5 Hz, and sham group datasets. Using this approach allowed us to illustrate several regional activation spots that are specific to a given stimulation frequency as well as consensual activation clusters (**Figure 3A**; statistical comparison shown in **Figure S2A**). While the ipsilateral LC shows similar activation in both tonic datasets (**Figure 3A**), BOLD signals in the contralateral LC selectively and significantly decreased during 5 Hz stimulation compared to sham and 3 Hz LC stimulation (**Figure 3A** and **Figure S2A-B**). Further, while both tonic stimulation conditions revealed activation of the thalamus compared to sham stimulation (**Figure 3A**), a significantly stronger activation was visible in the VPM during 5 Hz rather than 3 Hz stimulation (**Figure 3E** and **Figure S2A**). In contrast, BOLD responses within the ACC and ventral HP selectively increased upon 3 Hz (**Figure 3A**) but not sham and 5 Hz LC stimulation, resembling an inverted U-shape (**Figure 3B, D**). Direct comparison of BOLD activation maps also revealed a highly selective response to 5 Hz tonic LC firing in the amygdaloid complex (**Figure 3A**), where signals significantly decreased compared to sham and 3 Hz datasets (**Figure 3C**). These results suggest that across the tonic range, different intensities of LC firing affect activity in target brain regions selectively and non-linearly, presumably to tune cognitive performance and promote different levels of arousal ^26, 27, 44^.

**Figure 3.**
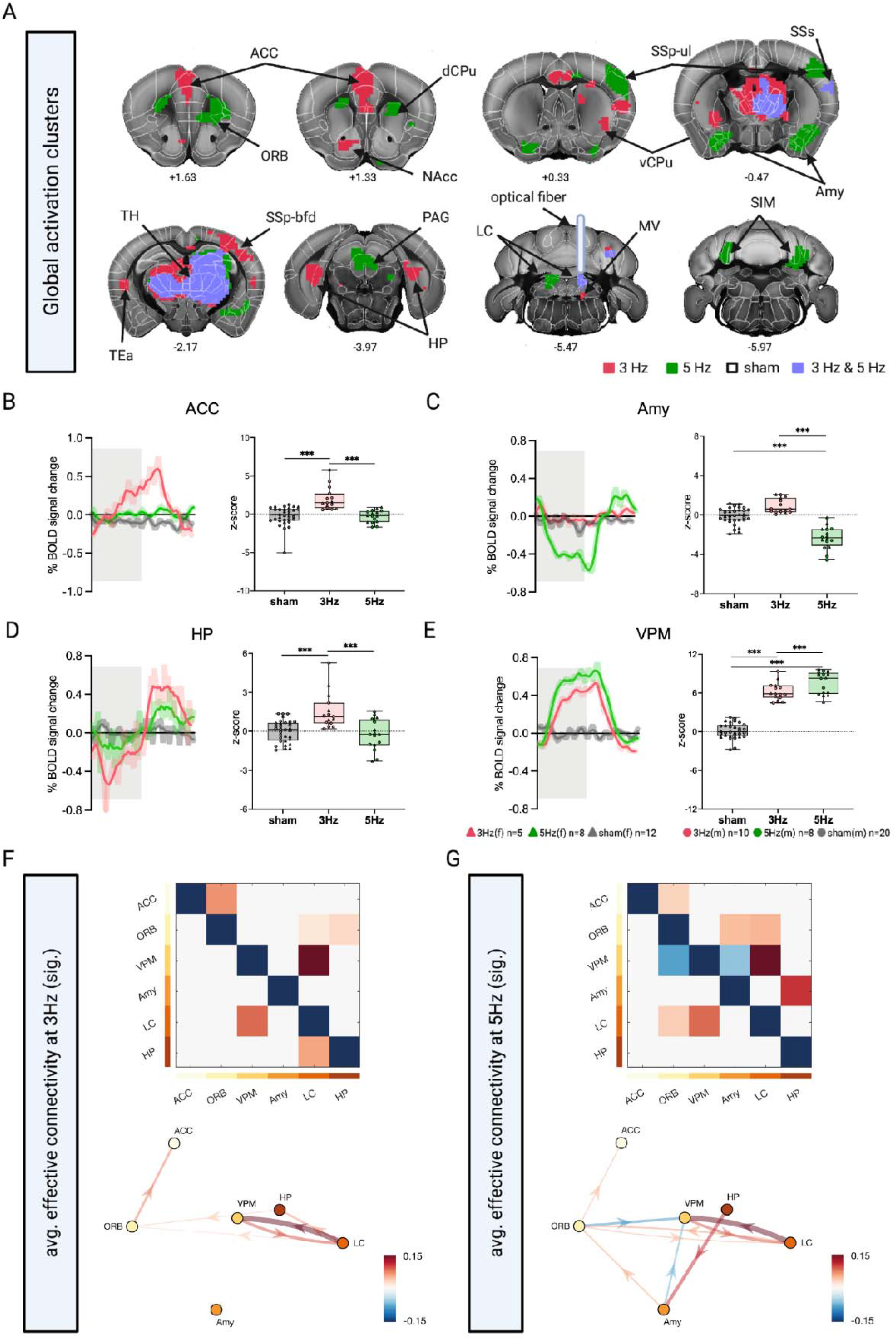
Tonic LC firing rates distinctly affect local BOLD activation patterns. **A** Selective and consensual activation clusters during 3Hz, 5Hz, and sham stimulation of LC noradrenergic neurons. Selective clusters of 3Hz datasets depicted in red, selective clusters of 5Hz datasets depicted in green, selective clusters of sham datasets depicted in white. Consensual clusters of 3Hz and 5Hz stimulations are depicted in lilac. **B** Mean time series of the ipsilateral ACC of sham, 3Hz tonic, and 5Hz tonic datasets. Z-scores of male (circle) and female (triangle) mice in the ipsilateral ACC significantly differed between stimulations (F(2,60)=12.927, p<.0005; MANOVA w/ LSD *post hoc* testing, Bonferroni correction). **C** Mean time series of the ipsilateral Amy of sham, 3Hz tonic, and 5Hz tonic datasets. Z-scores of male (circle) and female (triangle) mice in the ipsilateral Amy significantly differed between stimulations (F(2,60)=17.029, p<.0005). **D** Mean time series of the ipsilateral DG of sham, 3Hz tonic, and 5Hz tonic datasets. Z-scores of male (circle) and female (triangle) mice in the ipsilateral DG significantly differed between stimulations (F(2,60)=16.341, p<.0005). **E** Mean time series of the ipsilateral VPM of sham, 3Hz tonic, and 5Hz tonic datasets. Z-scores of male (circle) and female (triangle) mice in the ipsilateral VPM significantly differed between stimulations (F(2,60)=116.852, p<.0005). Averaged time-series of sham stimulation depicted in grey, 3Hz tonic stimulation in red, 5Hz tonic stimulation in green. Grey bar represents 30s laser stimulation blocks. Mean z-scores of sham, 3Hz and 5Hz stimulated mice depicted in grey, red and green, respectively. **F** Average connectivity pattern during the 3Hz stimulation condition, and **G** 5Hz stimulation condition. For both conditions, only significant (p<0.05, FDR-corrected for multiple comparisons) connections are shown, both as an adjacency matrix (*top*) as well as a network graph (*bottom*). ACC, anterior cingulate; Amy, amygdala; dCPu, dorsal caudate putamen; HP, hippocampus; ORB, orbitofrontal area; NAcc, nucleus accumbens; PAG, periaqueductal gray; SIM, simple lobule; SSp-bfd, primary somatosensory cortex, barrel field; SSp-ul, primary somatosensory cortex, upper limb; TEa, temporal association area; vCPu, ventral caudate putamen; VPM, ventral posteriomedial nucleus of thalamus. Plots represent ±SEM. ***p < 0.0005; *p < 0.05. N(sham)=32; n(3Hz)=15; n(5Hz)=16.

To test the effects of different tonic firing patterns on the directionality of the BOLD signal between specific brain areas (functional connections), we applied regression dynamic causal modelling (rDCM) ^45^ to the opto-fMRI data of 3 Hz and 5 Hz stimulation conditions. Specifically, using rDCM, we assessed the directed interactions (i.e., effective connectivity) within a 6-region network comprising LC and a selected set of its key target regions selected based on the regional activation spots that are stimulation frequency specific as well as shared clusters (see **Figure 3A**; ACC, ORB, VPM, Amy, HP). For the 3 Hz condition, we observed a strong driving input of the optogenetic stimulation to LC, consistent with the actual stimulation site (mean±SD: 0.13±0.09, one-sample t-test: t(14)=5.48; p<0.001). With regards to the endogenous connectivity (**Figure 3F**), LC exhibited strong efferent connections to ORB, VPM, and HP. Furthermore, significant effective connections were from VPM to LC, from ORB to ACC as well as from HP to ORB. For the 5 Hz condition, effective connectivity displayed similarities, but also differences compared to the 3 Hz condition. First, we again observed a strong driving input of the optogenetic stimulation to LC (mean±SD: 0.20±0.11, one-sample t-test: t(15)=7.44; p<0.001). Furthermore, strong reciprocal connections were again found between LC and VPM, as well as strong connections from LC to ORB and from ORB to ACC (**Figure 3G**). In contrast, for 5 Hz, we no longer observed a significant effect of LC to HP, nor a significant connection from HP to ORB. Instead, we observed a strong connection from HP to Amy, as well as pronounced inhibitory influences from ORB and Amy onto VPM. To explicitly test how the changes in effective connectivity differ between 3 Hz and 5 Hz stimulation, we performed an edgewise direct comparison (**Figure S2D**). This revealed a significant decrease of the connection from LC to HP during 5 Hz compared to 3 Hz, whereas a significant increase was observed for the connection from HP to Amy (p<0.05, FDR-corrected for multiple comparisons).

We further inspected the reconfigurations in the network across the two tonic stimulation conditions using graph-theoretical analyses. Specifically, we assessed the role of the above-mentioned regions with regards to their receiving or driving role in the network. Computing the asymmetry in node strength (i.e., the difference in strength between outgoing and incoming connections), we found a change in the role of HP from predominantly sink-like behaviour (i.e., stronger incoming than outgoing connections) during 3 Hz to driver-like behaviour (i.e., stronger outgoing than incoming connections) during 5 Hz tonic stimulation (**Figure S2E**). The opposite pattern could be observed for the Amy, although this did not survive multiple comparison correction (**Figure S2F**).

### Effects of burst-like LC activity on whole-brain activity

In addition to tonic activity, LC neurons have been shown to also fire in short bursts of higher frequency (∼10-25 Hz) ^13-15, 46^. Such firing can occur during focused task performance, but can also be externally triggered by unexpected, intense, or otherwise salient stimuli ^10, 14, 15^. Whether internally or externally generated, LC burst activity is thought to facilitate behavioural responses by increasing the likelihood of target neurons firing at precise, task-relevant times. These observations suggest that in addition to the spontaneous frequency of tonic LC firing, the temporal patterns of LC activity may have different downstream effects on whole-brain activity. To test this, we implemented an additional stimulation protocol, in which we drove LC activity with sub-second bursts of 15 Hz. This protocol was designed to mimic natural burst activity reported in LC neurons ^13-15^, and importantly, the total number of stimulation pulses delivered was matched between burst firing (15 Hz) and tonic 3 Hz stimulation (for visualisation, see **Figure 4A**). We found that 15 Hz LC stimulation evoked pupil dilations that were significantly larger than at 3 Hz tonic LC stimulation (mean±SD(3 Hz) = 13.89±6.37, mean±SD(15 Hz) = 17.02±6.13, paired t-test; t(23)=4.56, p<0.0001) (**Figure 4B-C**), suggesting that despite intensity-matched stimulation conditions, tonic and burst-like LC firing patterns differentially affect physiological responses. In line with this, NA levels measured by fibre photometry in the HP revealed greater NA release triggered by phasic compared to tonic LC stimulation (mean±SD(3 Hz) = 2.98±1.88, mean±SD(15 Hz) = 3.85±2.03, paired t-test; t(5)=3.57, p=0.016) (**Figure S3A**).

**Figure 4.**
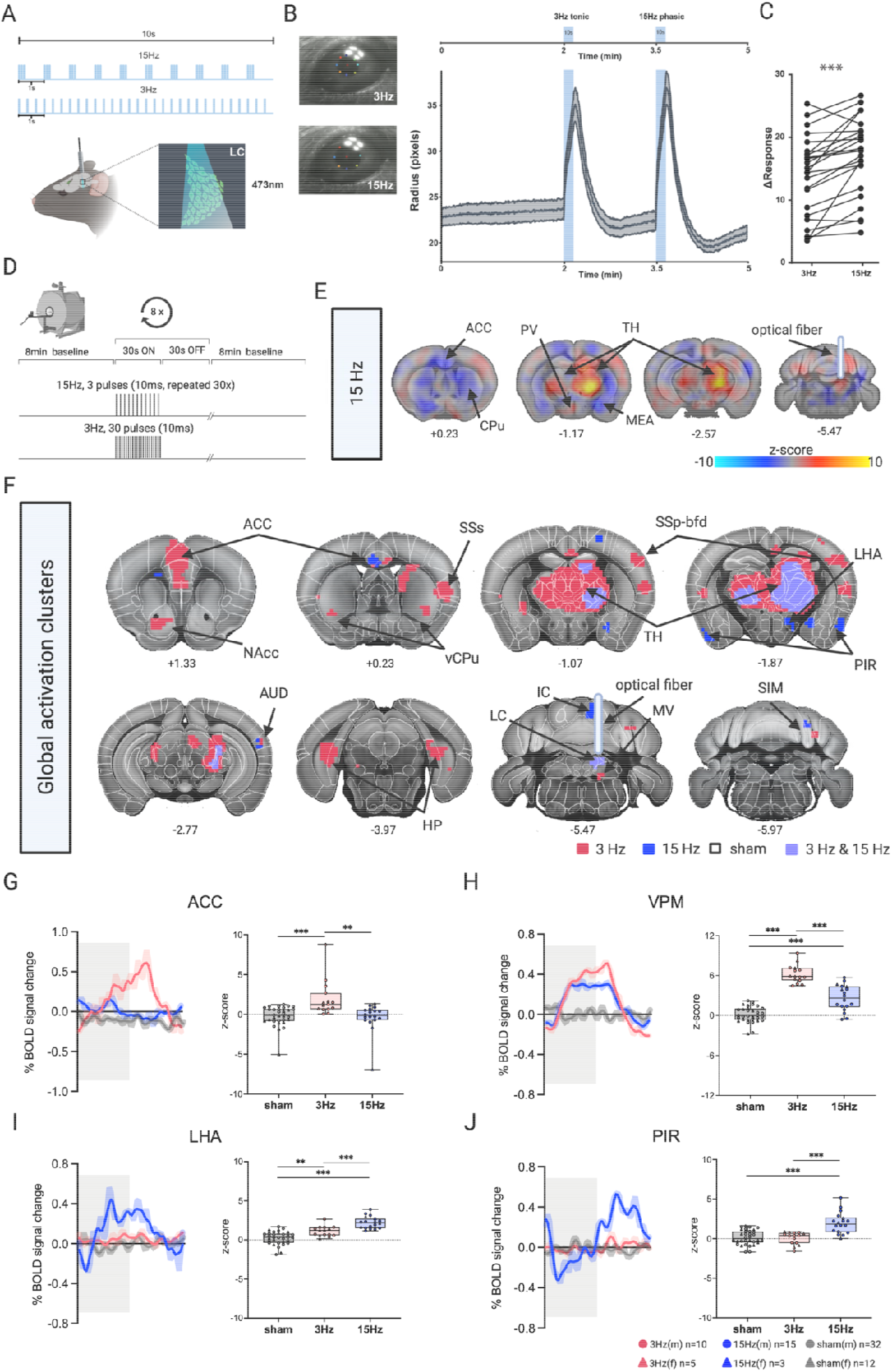
LC burst firing evokes distinct whole-brain activity patterns. **A** Schematic of 3Hz tonic and 15Hz burst blue laser-light stimulation for pupillometry. The protocol included a 2-minute baseline recording followed by 10 second stimulations: 1) 3Hz at 473nm and 2) 15 Hz at 473nm followed by a 90s post-stimulation baseline. **B** Pupil traces of the eye ipsilateral to the 3 Hz and 15 Hz stimulated LC. **C** Statistical comparison of the pupil response to tonic and burst-like LC stimulation revealed that at a 15Hz LC firing rate changes in pupil dilation were significantly greater than at3 Hz (mean±SD(3 Hz) = 13.89±6.37, mean±SD(15 Hz) = 17.02±6.13, paired t-test; t(23)=4.56, p<0.0001). **D** Unthresholded GLM z-stat activation map 15 Hz burst stimulation of LC noradrenergic neurons. **E** Selective and consensus activation clusters during 3Hz, 15Hz, and sham stimulation of LC noradrenergic neurons. Selective clusters of 3Hz datasets depicted in red, selective clusters of 15Hz datasets depicted in blue, selective clusters of sham datasets depicted in white. Consensual clusters of 3Hz tonic and 15Hz burst stimulation depicted in lilac. **F** Mean time series of the ipsilateral ACC of sham, 3Hz, and 15Hz datasets. Z-scores of male (circle) and female (triangle) mice in the ipsilateral ACC significantly differed between stimulations (F(2,62)=10.317, p<.0005; MANOVA w/ LSD *post hoc* testing, Bonferroni correction). **G** Mean time series of the ipsilateral VPM of sham, 3Hz, and 15Hz datasets. Z-scores of the ipsilateral VPM significantly differed between stimulations (F(2,62)=88.925, p<.0005). Z-scores of male (circle) and female (triangle) mice in the ipsilateral SUB significantly differed between stimulations (F(2,62)=25.563, p<.0005). **H** Mean time series of the ipsilateral LHA of sham, 3Hz, and 15Hz datasets. Z-scores of the ipsilateral LHA significantly differed between stimulations (F(2,62)=36.381, p<.0005). **I** Mean time series of the ipsilateral PIR of sham, 3Hz, and 15Hz datasets. Z-scores of male (circle) and female (triangle) mice in the ipsilateral PIR significantly differed between stimulations (F(2,62)=21.234, p<.0005). Mean z-scores of sham, 3Hz and 15Hz stimulated mice depicted in grey, red and blue, respectively. Time-series of sham stimulation depicted in grey, 3Hz tonic stimulation depicted in red, time-series of 15Hz tonic stimulation depicted in blue. Grey bar represents 30s laser stimulation blocks. ACC, anterior cingulate; Amy, amygdala; AUD, auditory area; DG, dentate gyrus; IC, inferior colliculus; LC, locus coeruleus; LHA, lateral hypothalamic area; MV, medial vestibular nucleus; NAcc, nucleus accumbens; PIR, piriform area; PV, periventricular hypothalamus; SIM, simple lobule; SSp-bfd, primary somatosensory cortex, barrel field area; SSs, supplementary somatosensory cortex; TH, thalamus; vCPu, ventral caudate putamen; VPM, ventral posteriomedial nucleus of thalamus. Plots represent ±SEM. ***p < 0.0005; **p<0.005; *p < 0.05. N(sham)=32; n(15Hz)=18; n(3Hz)=15.

Burst stimulation at 15 Hz during opto-fMRI recordings (**Figure 4D**) evoked increased BOLD responses in the periventricular hypothalamic nucleus (PV) and the ipsilateral thalamus, but negative responses in the bilateral CPu, posterior ACC and in the ipsilateral medial amygdalar nucleus (MEA) (**Figure 4E**). When illustrating similarities and differences between BOLD activity patterns of 3 Hz tonic vs 15 Hz burst datasets, we saw no differences in the targeted LC, suggesting that total laser stimulation was successfully intensity-matched between protocols (**Figure 4F** and **Figure S3C;** statistical comparison in **Figure S3B**). Like the 3 Hz tonic time-course, burst stimulation provoked a delayed BOLD response in the targeted LC, initially presenting with a dip and then gradually increasing (**Figure S3C**). Further, we found that the ACC and ventral HP was activated more strongly upon tonic rather than burst LC stimulation (**Figure 4G-H** and **Figure S3B**). Signals in the thalamus showed clusters selective towards 3 Hz tonic stimulation as well as consensually activated areas (**Figure 4F**). In contrast, the ipsilateral lateral hypothalamic area (LHA), the bilateral piriform area (PIR) (**Figure 4F, I-J**) as well as the inferior colliculus (IC) (**Figure S3B, D**) showed greater BOLD signal increases upon 15 Hz burst LC stimulation. This suggests that activity within these brain structures is selectively modulated by an LC burst rather than tonic firing pattern.

As before, we utilised rDCM to assess the effective connectivity during the 15 Hz burst stimulation condition within the same 6-regions network analysed above and again observed strong reciprocal connectivity between LC and VPM (**Figure S3E**). Furthermore, reciprocal connections were present between LC and ORB, as well as efferent connections from ORB to HP and from HP to Amy. While this pattern shows some differences to the pattern observed during 3 Hz, a direct edgewise comparison between the 3 Hz tonic and 15 Hz burst stimulation condition did not reveal any significant differences after multiple comparison correction.

### LC tonic and burst firing patterns differentially reshape cortical energy landscapes

Spatially coordinated, temporally sustained patterns of neural activity give rise to brain states that can change discretely or continuously, with or without external stimuli ^47^. As a structure that lies at the intersection of cortical and subcortical structures, the thalamus is critically involved in such systems-level dynamics ^48-51^. Interestingly, our analyses revealed robust BOLD signal responses in the VPM thalamic relay nucleus during both tonic and burst-like LC stimulation patterns (**Figure 3A, E** and **Figure 4E, G**) as well as strong effective connectivity from LC to VPM in all three stimulation conditions. This suggests that LC-activation induced a modulation of brain states. Earlier work based on human neuroimaging data had suggested that LC activation would facilitate such dynamic changes by lowering energy barriers between states ^31^. Our opto-fMRI data enabled us to directly test this hypothesis, by quantifying the distinct effects of optogenetically-evoked tonic and burst-like LC firing on cortical and thalamic dynamics using both topological and energy landscape analyses. Briefly, this latter approach estimates the energy required to reach a given brain state (*E*_*state*_) defined as the natural logarithm of the state’s inverse probability (*P*_*state*_): □_□□□□□_ = ln(*1*/□_□□□□□_) (**Figure 5A, left**). A state of low energy corresponds to a high probability state and vice versa, and we hypothesized that LC activation would lower the energy barrier between states (**Figure 5A, right**) ^31^. It should be noted that this definition of energy differs from metabolic energy, recapitulating its use in statistical mechanics. This technique allowed us to quantify the amount of energy that it would take to change the brain state (BOLD activation) by a certain amount (defined by the mean squared displacement, MSD) within a given temporal window (repetition time, TR).

**Figure 5.**
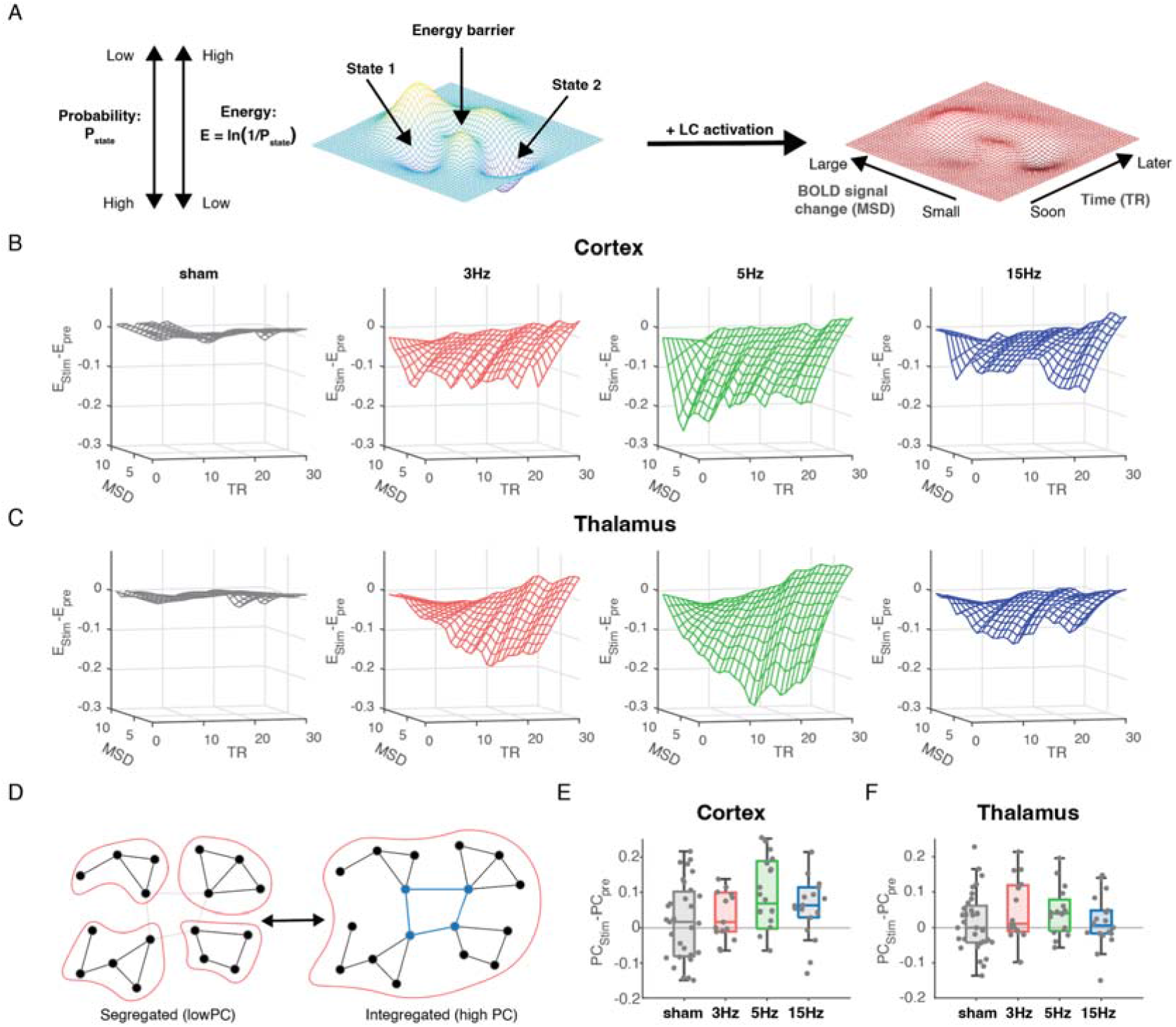
Tonic and burst-like LC firing patterns differentially affect cortical and thalamic dynamical topography and network topology. **A** BOLD dynamical energy landscapes quantify the likelihood of observing a brain-state transition as a function of BOLD change quantified by the mean-squared displacement (MSD) and time (TR). We hypothesised LC activation would reduce high energy barriers facilitating novel BOLD changes. Mean Cortical **(B)** and thalamic **(C)** BOLD energy landscape as a function of mean-squared displacement (MSD) and TR during sham (black), 3Hz tonic (red), 5Hz tonic (green), and 15Hz burst (blue) LC stimulation calculated across all subjects, first corrected for pre-stimulation baseline. **D** BOLD network changes were quantified by the participation coefficient. Mean Cortical **(E)** and thalamic **(F)** participation coefficients during LC stimulation in sham, 3Hz tonic (red), 5Hz tonic (green), and 15Hz burst (blue) datasets. Corrected for pre-stimulation baseline. N(sham)=32; n(15Hz)=18; n(3Hz)=15; n(5Hz)=16.

In support of our hypothesis, LC stimulation significantly decreased the transition energy for large BOLD MSD in both cortical and thalamic parcels relative to sham (all p<0.05, paired t-test). The reduction in energy was pronounced for tonic stimulation with the largest effects observed in cortex (**Figure 5B**) and thalamus (**Figure 5C**) following 5 Hz LC stimulation. Thalamic energy landscapes following tonic stimulation demonstrated a temporally lagged reduction, whereas cortical modification was constant. Burst stimulation led to a more subtle reduction in energy landscape and a rapid return to pre-stimulation dynamics - i.e., no difference between pre- and post-stimulation energy landscapes (p>0.05, paired t-test). In the post-stimulation period, in contrast, tonic stimulation led to a large increase in energy for all MSD and TR (p<0.01, paired t-test; **Figure S4A, C**). Thus, the pronounced reduction of energy for large BOLD changes *during* tonic stimulation was followed with a rebound ‘exhaustion’ or stabilisation of cortical and thalamic BOLD activity *after* the end of LC stimulation.

In addition, our data showed a significant increase in mean cortical participation coefficient (PC), a measure of network integration (**Figure 5D**), following 5 and 15 Hz LC stimulation (**Figure 5E**, p<0.05, MANOVA Bonferroni correction). In contrast, significant changes were only observed following tonic stimulation in the thalamus (**Figure 5F**, p<0.05 3 & 5 Hz). Previous research has linked noradrenergic activity to enhanced system integration^1, 52, 53^, arguing that the LC-NA system is uniquely positioned to create a more integrated network topology^2, 31^: due to its highly diffuse neuroanatomy, NA projections from the LC typically cross multiple boundaries along the neuraxis, allowing activity to be coordinated between otherwise segregated regions^54^. Our findings show that LC stimulation has an influence on the systems-level configuration of brain networks *in vivo*, with varying cortical and thalamic responses depending on the frequency of LC activation. This is noteworthy because it is the first empirical evidence of how LC-NA firing patterns and frequency causally impact network dynamics.

## Discussion

Neuromodulation is one of the key processes that endows the brain’s relatively static structural architecture with flexibility, making it possible to support malleable neural dynamics required for adaptive behaviour ^21, 22^. As part of the neuromodulatory ascending arousal system, the LC-NA system is well-placed to mediate this role ^1, 3^. Previous animal and human studies have hinted that high-tonic LC activity can rapidly reconfigure functional large-scale network architecture to facilitate coordination between otherwise segregated regions ^1, 2, 23, 32^. Here, we address the question whether physiologically relevant variations in the tonic firing frequency of the LC, as well as modulation of firing patterns, differentially impact brain dynamics. We used a combined optogenetic-fMRI approach to test levels of tonic as well as burst LC firing and visualise their effects on brain-wide activity in the mouse. Across the tonic range, we report that LC noradrenergic stimulation evokes forebrain responses in a causal and non-linear manner, with distinct modulatory effects in downstream target structures. This is important, because it emphasizes the need to carefully consider the stimulation paradigms when assessing the impact of LC activity on brain function ^55^. In addition, we observe that the firing pattern differentially impacts the functional integration (i.e., effective connectivity) amongst LC and key downstream target regions. We also provide causal proof that increasing the tonic firing frequency of the LC directly flattens the energy landscape of the brain, thus facilitating state transitions. These findings are important because they offer an explanation how strong LC activity can trigger a network reset in response to rapidly changing environmental demands ^5, 32, 33^.

LC neurons exhibit baseline firing within a range of low tonic to high tonic rates that is linearly related to NA efflux in target tissues ^24, 56^. However, across tonic discharge rates, the modulatory effects of LC on vigilance are thought to be conveyed in a non-linear fashion, following an inverted U-shaped curve ^37, 43, 57^: optimal cognitive performance and focused attention relates to intermediate levels of tonic LC-NA activity ^11, 36, 43^, while higher tonic LC-NA activity can lead to increased distractibility and even anxiety-like behaviours ^11, 38, 58, 59^. We theorised that such non-linearities similarly exist at the mesoscale, with different LC firing frequencies evoking specific activation patterns in distinct target brain regions. Indeed, our results support this prediction by showing that 3 Hz vs 5 Hz tonic stimulation rates evoke specific BOLD activity profiles in different brain structures. For example, we discovered dissociable effects of the two different tonic LC stimulation intensities in prefrontal cortical areas: while the mPFC responded exclusively to a 3 Hz frequency, the ORB responded to the 5 Hz but not the lower tonic stimulations. The mPFC is not only critically involved in several cognitive and executive functions ^60^, but reciprocal connections with the LC have been shown to contribute to volitional control of waking as well as a variety of task-specific behaviours requiring vigilance and arousal ^46, 61^. Arnsten and colleagues have proposed that NA release in prefrontal areas could act as a ‘switch’ to either selectively enhance or suppress such functions via differential activation of adrenoceptors ^62^. In line with this, moderate release of NA upon our 3 Hz LC stimulation might have acted on high-affinity α2 adrenoceptors to selectively engage and improve the regulation of medial prefrontal target regions ^63, 64^. In contrast, higher NA release at 5 Hz might have led to dampening of mPFC function via co-activation of lower-affinity α1 and β receptors ^65-67^, and instead recruited orbitofrontal functions related to reorientation and exploration ^68, 69^. Of note, it is currently also unclear how different amounts of NA release might impact different cell types, due to differences in adrenoceptor expression. As such, these findings suggest that in addition to being innervated by distinct subsets of LC neurons ^60, 70^, prefrontal function might further be dynamically regulated via different tonic LC firing rates and associated NA release levels. Instead, BOLD signals in the PAG and amygdala selectively responded to 5 Hz stimulation only. In the latter, LC noradrenergic projections have been shown to be critically involved in the adaptive tuning of emotional responding ^59^. More specifically, optogenetic manipulation of LC-NA inputs into the basolateral amygdala at a 5 Hz tonic frequency has been shown to be sufficient to produce conditioned aversion and induce anxiety-like behaviours ^38, 59^. In contrast, BOLD responses in the thalamic VPM region increased linearly with increasing tonic firing rate. Importantly, LC excitation or exogenous application of NA onto single cells has been shown to evoke both inverted-U modulatory profiles and dose-dependent response curves within the thalamus ^71, 72^. Tonic LC activation could thus differentially impact the responsiveness of thalamic neurons, generating a state of readiness to appropriately encode and respond to salient stimulus inputs, maximising adaptive fitness. Indeed, a recent study showed enhanced sensory processing in the thalamic VPM nucleus and increased perceptual performance in a tactile discrimination task during tonic LC stimulation at 5 Hz ^73^. In support of these findings, our results suggest that across the tonic firing range brain activity is non-linearly and dynamically altered, possibly giving rise to LC-mediated shifts in levels of arousal, cognitive performance, and sensory processing.

Intense or otherwise motivationally salient stimuli have been shown to evoke burst-like LC activity ^15^. While it is thought that tonic LC background activity continually modulates arousal, burst firing is believed to serve as a ‘wakeup call’ to facilitate increased task engagement and performance by differentially affecting brain activity ^23, 25^. In line with this idea, we show that burst stimulation of LC noradrenergic neurons not only selectively modulates brain regions involved in sensory perception and attention like the PIR and IC, but also differentially affects cortical and thalamic dynamical topography. In particular, following burst stimulation cortical and thalamic dynamics quickly recovered to pre-stimulation dynamics. This alludes to the possibility that through LC bursts evoking a rapid but transient drop in the energetic landscape, only few (yet selective) network changes are favoured. In contrast, peaks in tonic LC activity generally led to a prolonged deepening of the energy landscape – possibly allowing brain regions to radically shift their connectivity profiles, consistent with the network-reset idea of LC activity ^5^. Consequently, the pronounced post-tonic-stimulation rebound suppressing large changes in BOLD hints at a state of ‘exhaustion’ following energetically demanding global network reconfigurations. In accordance with the concept of the LC-NA system acting similar to a catalyst in a chemical reaction, our results suggest that LC activity patterns lower the energetic demand for brain state transitions with increasing firing frequency, conferring adaptive benefits across a spectrum ^5^. Additionaly, these results provide confirmatory, causal evidence that the LC can alter spatiotemporal brain network dynamics in an intensity-sensitive fashion.

### Limitations

We acknowledge that unilateral and unimodal optogenetic stimulation sustained over 30 seconds does not represent a physiological mode of operation of the LC-NA system. In fact, the exact firing patterns of LC neurons in response to various environmental challenges such as stressful stimuli remain unknown, due to difficulties in recording large numbers of LC neurons, particularly in freely moving mice. The highest density recordings in anesthetized rats to date suggest highly variable firing of LC neurons, where only a fraction (15%) display synchronous firing associated with phasic discharge, even in response to a noxious foot shock ^74^. However, it is conceivable that this number might increase in vivo in a scenario where multimodal stress signals converge and accumulate over minutes or hours. In vivo, firing in a novel environment showed prominent burst activity in the range of 15-28 Hz sustained over several minutes ^13^. Thus, one challenge for future work will be to decipher the natural firing properties of LC neurons over longer time scales and in response to various situational demands (sustained attention, salient stimuli, novelty, stress), and subsequently refine stimulation protocols to closely resemble (or even replay) naturally occurring LC activity patterns. In addition, we cannot exclude potential influences of the contralateral hemisphere on our fMRI signals. To this point, decreased activity in the contralateral LC during 5 Hz stimulation might have contributed to shaping the distinct whole-brain activation profiles of the higher tonic dataset. Finally, our work is not able to address the modular organization of the LC into cellular ensembles ^75, 76^, as our stimulation will uniformly drive LC activity irrespective of afferent and efferent projections.

## Conclusions

In this work, we discovered that LC firing patterns evoke distinct and often non-linear changes in brain activity and network reconfigurations at the systems level. We show that tonic and burst-like LC stimulation causes brain-wide activation profiles that are specific not only to the firing frequency but also the dominating firing pattern. We utilise generative modelling to identify changes in effective connectivity that might be underlying these frequency-specific activation profiles. Additionally, we demonstrate how both tonic and burst-like LC firing patterns are related to dynamic modulation of whole-brain low-dimensional energy landscapes and network topology, presenting a view of the LC-NA system in gating brain state transition. Together, our results provide novel anatomical insight into how the LC-NA system continuously and adaptively modulates brain dynamics to support cognitive functions and ongoing behaviours.

## Acknowledgements

We thank Gabriel Wainstein and Eli J. Müller for the useful discussion and brilliant intuitions on the role of LC on brain states. We also thank Jean-Charles Paterna from the Viral Vector Facility (VVF) of the Neuroscience Center Zurich, a joint competence center of ETH Zurich and the University of Zurich, for producing viral vectors and viral vector plasmids. C.G and V.Z are supported by the research grant ETH 062-18 and the Swiss National Science Foundation (SNSF) AMBIZIONE (PZ00P3_173984/1) and ECCELLENZA (PCEFP3_203005). J.B. is supported by the ETH Zurich, ETH Project Grant ETH-20 19-1, SNSF Grant 310030_172889 and the Swiss 3R Competence Center. J.S acknowledges support by the National Health and Medical Research Council (GNT1193857). Graphics created by C.G and S.N.D. using BioRender.com.

## Author Contributions

Conceptualization, C.G, S.N.D, M.P, B.M, S.F, G.W, J.M.S, J.B, and V.Z; Methodology, C.G, S.N.D, M.P, B.M, S.F, G.W, M.C, J.M.S, and V.Z; Investigation, C.G, S.N.D, M.P, B.M, S.F, J.M.S, and V.Z; Writing – Original Draft, C.G, S.N.D, M.P, B.M, S.F, J.M.S, and V.Z; Writing – Review & Editing, C.G, S.N.D, M.P, B.M, S.F, M.C, J.M.S, D.R, N.W, J.B, and V.Z; Funding Acquisition, V.Z, J.S, N.W, D.R, T.P. and J.B; Supervision, D.R, N.W., J.M.S, J.B, and V.Z.

## Declarations of Interest

The authors declare no competing interests.

## Methods

### Animals

All animal procedures were conducted in accordance with the Swiss federal guidelines for the use of animals in research and approved by the Cantonal Veterinary Office of Zurich. Heterozygous C57BL/6-Tg(Dbh-iCre)1Gsc (DBH-iCre) mice ^77^ were kept in standard housing on a 12h light/dark cycle, with food and water provided *ad libitum*. A total of 87 fMRI scans were acquired with adult, heterozygous male (n=22) and female (n=7) DBH-iCre mice. Six fMRI scans were excluded due to fMRI coil-related artefacts, leaving a final dataset of 81 scans.

### Stereotaxic surgery

Stereotaxic surgery was performed on 2 to 3 months old DBH-iCre mice under 2% isoflurane anaesthesia with a subcutaneous dose of 5 mg/kg Meloxicam and a local anaesthetic (Emla cream; 5% lidocaine, 5% prilocaine). Animals were placed into a stereotaxic apparatus and their skulls exposed. Bregma was located and the skull placement corrected for tilt and scaling. For virus delivery and optical fibre implantation in the right locus coeruleus, a small hole was drilled at AP -5.4 mm and ML -0.9 mm, relative to bregma. Mice were then injected unilaterally (Coordinates: AP -5.4 mm, ML -0.9 mm, DV -3.8 mm), with 1µl of an AAV construct carrying the optogenetic actuator ChR2 or ChrimsonR (ssAAV-5/2-hEF1α-dlox-hChR2(H134R)_EYFP(rev)-dlox-WPRE-hGHp(A) or ssAAV-5/2-hEF1α/hTLV1-dlox-ChrimsonR_tdTomato(rev)-dlox-WPRE-bGHp(A); Viral Vector Facility (VVF), Neuroscience Center Zurich) using a pneumatic injector (Narishige, IM-11-2) and calibrated microcapillaries (Sigma-Aldrich, P0549). For fibre photometry recordings in the hippocampus, mice were additionally injected with 0.2µl of the genetically encoded NA sensor GRAB_NE1m_ (ssAAV-9/2-hSyn1-GRAB(NE1m)-WPRE-hGHp(A); VVF, Neuroscience Center Zurich; coordinates: AP -3.2 mm, ML -3.3 mm, DV -3.8 mm) ^41^. Subsequently, an optical fibre was implanted at 200μm superior to the injection coordinates (for fMRI: low profile, 90° 200μm, NA=0.66; Doric Lenses, Canada, for fibre photometry, 200µm, NA=0.37; Neurophotometrics, USA). Optical fibres were glued to the skull using a bonding agent (Etch glue, Heraeus Kulzer GmbH) and a UV-curable dental composite (Permaplast, LH Flow; M+W Dental, Germany) and stitches were used as required. The health of the animals was evaluated by post-operative checks over the course of 3 consecutive days and 5 mg/kg Meloxicam administered subcutaneously if needed.

### Pupil recordings

For pupil recordings, a Raspberry Pi NoIR Camera Module V2 night vision camera, an infrared light source (Pi Supply Bright Pi - Bright White and IR camera light for Raspberry Pi) and a Raspberry Pi 3 Model B (Raspberry Pi Foundation, UK) was used. Experimental procedures in all animals followed guidelines as detailed in ^34^. In short, mice were anesthetized in an induction chamber with isoflurane in a 1:4 O2 to air mixture (4% induction, 2% maintenance). The eye ipsilateral to the stimulated LC (right eye) was recorded and a 2-minute baseline recording preceded the different stimulation paradigms. Stimulation patterns lasted 10s and included 1) a 5 Hz sham stimulation (635nm, 10ms pulse width, 10mW laser power), 2) a 3 Hz (473nm, 10ms pulse width, 10mW laser power), 3) a 5 Hz (473nm, 10ms pulse width, 10mW laser power), and 4) a 15 Hz burst stimulation (3 pulses/s, 473nm, 10ms pulse width, 10mW laser power). All stimulations were followed by 90 sec no laser light delivery.

### Fibre photometry

The green fluorescence signal from NA sensor GRAB_NE1m_ was recorded using a commercially available photometry system (Neurophotometrics, Model FP3002) controlled via the open-source software Bonsai (2.6.2 version). Throughout the recording session mice were lightly anesthetized (4% isoflurane during induction, 1.5-2% during maintenance) and the fibre implanted in the mouse brain was attached to a pre-bleached recording patch cord (200 μm, 0.39 numerical aperture; Doric Lenses). Two LEDs were used to deliver interleaved excitation light: a 470 nm LED for recording NA – dependent fluorescent signal (F^470^) and a 415 nm LED for NA – independent control fluorescent signals (F^415^). Recording rate was set at 120 Hz for both LEDs allowing 60 Hz for each channel individually. Excitation power at the fibre tip was set to 25-35 μW. During the same photometry recording session, the LC was stimulated using 3 Hz tonic, 5 Hz tonic, and 15 Hz burst laser pulses in a randomized order (635 nm laser (CNI laser) at 5 mW output power).

Analysis of the raw photometry data was performed using a custom-written MATLAB script. First, to filter high frequency noise (above 1 Hz), the lowpass filter function was applied to both recorded signals (F^470^ and F^415^). Next, to correct for photobleaching of fluorescent signal, the baseline fluorescence F^415^_baseline fit_ was calculated as a linear fit applied to the filtered F^415^ to F^470^ signals during the baseline 5 s window preceding each LC stimulation. Finally, the signal of the NA sensor was expressed as a percentage change in fluorescence: ΔF/F = 100*(F^470^(t) – F^415^_baseline fit_(t))/F^415^_baseline fit_(t), where F^470^(t) signifies the filtered fluorescence value at each time point *t* across the recording and F^415^_baseline fit_(t) denotes the value of the fitted 415 nm signal at the time point *t*. The final ΔF/F signal was smoothed with a 100 - points moving mean filter.

### Opto-fMRI recording

#### Animal preparation

For fMRI scans, mice were anesthetized in a gas chamber for 4 minutes with 4% isoflurane in 1:4 O2 to air mixture. Animals were endotracheally intubated and the tail vein cannulated while being kept under anaesthesia with 2% isoflurane. During preparation, animal temperature was kept at 35 °C using a heating pad (Harvard Apparatus, USA). Once intubated and cannulated, mice were head-fixed with earbars and connected to a small animal ventilator (CWE, Ardmore, USA) on an MRI-compatible support. Ventilation was set to 80 breaths per minute, with 1.8 ml/min flow with isoflurane at 2%. A bolus injection of a muscle relaxant (pancuronium bromide, 0.25mg/kg) was delivered via the cannulated vein and isoflurane was reduced to 1.5%. A fibre optic patch cord (Thorlabs, USA) connected to a custom-made DPSS laser (CNI laser, China) was tethered to the optical fibre implant via an MRI compatible connector. Continuous infusion of pancuronium bromide (0.25mg/kg/h) started five minutes after the initial bolus injection. Isoflurane was reduced to 1.1%. A hot water-circulation bed kept the temperature of the animal constant throughout the entire measurement (35 °C). Additionally, body temperature was monitored using a rectal thermometer probe. After collection of functional and anatomical fMRI scans, continuous injection and isoflurane flow was stopped. Animals remained connected to the ventilator until independent breathing could be assured and then transferred to a heating chamber for further recovery.

#### Data acquisition

Data were acquired in a 7T Bruker BioSpec scanner equipped with a Pharmascan magnet and a high SNR dedicated mouse brain cryogenic coil (Bruker BioSpin AG, Fällanden, Switzerland). Standard adjustments included the calibration of the reference frequency power and the shim gradients using MapShim (Paravision v6.1). For anatomical assessment, a T1-weighted image was acquired via a FLASH sequence with an in-plane resolution of 0.05 × 0.02 mm^2^, an echo time (TE) of 3.51 ms and a repetition time (TR) of 522 ms. For functional scans, a standard gradient-echo echo-planar imaging sequence (GE-EPI, repetition time TR = 1 s, echo time TE = 15ms, in-plane resolution RES = 0.22 × 0.2mm^2^, number of slice NS = 20, slice thickness ST = 0.4 mm, slice gap = 0.1mm) was applied to acquire 1440 volumes in 24 min.

#### Optogenetic stimulation

Frequency-dependent whole-brain activity of ChR2-EYFP expressing DBH-iCre mice was recorded in three separate functional scans. Over the course of four weeks, mice were randomly assigned 3 Hz, 5 Hz, 15 Hz, and sham optogenetic stimulation protocols (pulse width 10 ms). Functional scans comprised an eight-minute pre-stimulation baseline, an eight-minute optogenetic stimulation phase, and an eight-minute post-stimulation baseline. For 3 Hz and 5 Hz tonic optogenetic stimulation, trains of 473 nm laser pulses were delivered for 30 sec at 10 mW laser power above the targeted site, followed by 30 sec of no laser light delivery. This stimulation paradigm was repeated over the course of eight minutes. For LC burst stimulation, a burst of 3 pulses at 15 Hz (10mW and 473nm) was delivered every second for 30 sec, followed by 30 sec of no laser light delivery. The stimulation was repeated over the course of eight minutes. For sham optogenetic stimulation, trains of 635 nm laser pulses were delivered at 3-5 Hz for 30 sec and 10 mW laser power above the targeted site, followed by 30 sec of no laser light delivery. The stimulation was repeated over the course of eight minutes. Mice were randomly subjected to any of the four optogenetic stimulation protocols.

### Immunohistochemistry

The hindbrain including the locus coeruleus was fixed in 4% PFA for 2 hours, cryoprotected in a sucrose solution and frozen in mounting medium. The hindbrain was then cut into 40µm sections and stained in primary antibody solution containing 0.2% Triton X-100, and 2% normal goat serum in PBS at 4°C under continuous agitation over 2 nights. Afterwards, sections were washed 3 times in PBS for 10 minutes per cycle and then transferred into secondary antibody solution containing 2% normal goat serum in PBS. After 3 more PBS washes, the sections were mounted onto glass slides (Menzel-Glaser SUPERFROST PLUS, Thermo Scientific), air-dried and cover-slipped with Dako fluorescence mounting medium (Agilent Technologies). Primary antibodies included mouse anti-TH (22941, Immunostar, 1:1000), chicken anti-GFP (ab13970, Abcam. 1:1000), and rabbit anti-cFOS (226 003, Synaptic Systems,1:5000). Secondary antibodies included donkey anti-mouse Alexa 647 (A-31571, Thermo Fisher Scientific, 1:300), goat anti-chicken Alexa Fluor 488 (A-11039, Thermo Fischer Scientific, 1:300), and goat anti-rabbit Alexa 546 (A11035, Thermo Fisher Scientific, 1:300). Microscopy images were acquired in a confocal laser-scanning microscope (CLSM 880, Carl Zeiss AG, Germany). Images of LC were acquired using a 10x or 20x objective.

### Quantification and Statistical Analysis

Statistical details for every experiment are provided in the figure legends, where “n” represents the number of animals per group. Statistical significance was defined as p < 0.05.

#### Quantification

Pupil diameter was measured using the motion tracking software DeepLabCut ^78, 79^. The web-based Pupillometry App reported in ^34^ was used for visualization and analysis. Measurements after 3 Hz, 5 Hz, and 15 Hz optogenetic stimulation were normalized to the 10s baseline preceding laser light delivery and statistically analysed as in ^34^.

#### fMRI data analysis

*GLM statistical mapping*. Preprocessing and functional data analysis was carried out using FSL FEAT (version 5.92, www.fmrib.ox.ac.uk/fsl) and in-house Matlab scripts. Pre-statistical processing included the following steps: Pre-processing of the BOLD data included discarding the first ten measurements to achieve steady-state excitation, high-pass filtering (with a cut-off of 90s), motion correction using MCFLIRT, spatial smoothing using a Gaussian kernel of FWHM 0.4 mm and interleaved slice-timing correction. To account for potential alignment artefacts due to the implanted optical fibre, two study-specific templates based on all mean-EPIs and T1-weighted anatomical images were created using Advanced Normalization Tools (ANTs, http://stnava.github.io/ANTs/). Registration was first carried out to the respective T1-weighted image and then to the study-specific standard space template, using FLIRT and FNIRT. Time series statistical analysis was carried out using FILM with local autocorrelation correction. Importantly, the BOLD response evoked by LC stimulation deviates from a standard response showing an initial dip followed by a delayed increase most likely because NA release affects not only neural but also vasomotor activity. Thus, first-level (time series) parameter estimates were computed using a mass-univariate general linear model based on a custom hemodynamic response function (FSL FLOBS) and its temporal derivatives sampled during increased neuromodulatory activity ^42^, reflecting the delayed increase in BOLD activity during optogenetic LC-NA stimulation. With standard motion parameters (MCFLIRT) applied, group-level analysis was performed using FMRIB’s Local Analysis of Mixed Effects (FLAME). An averaged whole-brain map was created for the LC laser stimulation ON versus laser stimulation OFF contrast. In a third step, whole-brain differences between groups were tested (group contrast activation). Z-statistic images were thresholded using clusters determined by z□>□3.1, and a familywise error–corrected cluster significance threshold of p□<□0.05 was applied to the suprathreshold clusters.

For visualization purposes, averaged group and group contrast activation maps were normalized to a high-resolution Allen Brain Institute (ABI) anatomical atlas using affine and non-linear greedy transformations (ANTs).

##### Region of interest analysis

Anatomical regions-of-interest (ROIs) were selected and manually defined based on significant clusters obtained from between-group statistical maps for 3 Hz, 5 Hz, and 15 Hz optogenetic LC stimulation. A total of 14 ROIs were identified and anatomically labelled: anterior cingulate (ACC), amygdala (Amy), striatum (CPu), inferior colliculus (IC), locus coeruleus ipsilateral (LC), locus coeruleus contralateral (LC contralateral), somatosensory cortex (SSCtx), simple lobule (SIM), lateral hypothalamic area (LHA), periaqueductal gray (PAG), hippocampus (HP), piriform cortex (PIR), subiculum (SUB), and ventral posteriomedial nucleus of the thalamus (VPM). Mean time-series for selected anatomical ROIs were extracted, baseline-corrected, and their percent BOLD signal changes calculated. For BOLD magnitude comparison, mean z-score values were extracted for selected anatomical ROIs from single-subject BOLD activation maps.

##### Selective activity cluster analysis

Voxels that showed a significant and selective response in sham, 3 Hz tonic, 5 Hz tonic or 15 Hz burst datasets were selected for illustration purposes using an in-house Matlab script. Briefly, for identification of selective voxels, group-level zstat activation maps of each stimulation condition were thresholded at 3.1 and iteratively compared to the other stimulation datasets. Only voxels that met the imposed threshold in one dataset, but not the others were identified as selective activity clusters.

#### Regression dynamic causal modelling

The following six regions of interest (ROIs) were selected for the regression dynamic causal modelling (rDCM) analysis: (i) LC, (ii) ACC, (iii) ORB, (iv) VPM, (v) Amy, and (vi) HP. Extracted BOLD signal time courses from those regions (see above) entered effective (directed) connectivity analyses using rDCM. Specifically, we utilised the open-source rDCM toolbox, which is freely available as part of the TAPAS software package (www.translationalneuromodeling.org/tapas) ^80^. In brief, rDCM is a novel variant of DCM for fMRI ^81^ that renders effective connectivity analyses highly efficient by reformulating the differential equations of a classical DCM in the time domain into an efficiently solvable Bayesian linear regression in the frequency domain ^45, 82^.

Here, we used the original rDCM implementation that assumes a fixed network architecture where the presence or absence of connections is defined *a priori*. To minimise the influence of our prior assumptions about the network architecture on the rDCM results, we assumed a fully connected (all-to-all) network where all regions are reciprocally connected to each other. Starting from this fully connected network, during model inversion, the data (i.e., likelihood) were then able to drive posterior parameter estimates away from zero (i.e., the prior mean) for those connections where sufficient evidence for an effect was present. Conversely, for the driving input, we incorporated our knowledge about the actual optogenetic stimulation site into the model by setting a driving input to LC only, while all other regions received no driving input.

Model inversion was then achieved using Variational Bayes and was performed under the standard settings of the rDCM toolbox (e.g., priors). After model inversion, we inspected the posterior parameter estimates and significance of these parameter estimates was assessed using edgewise (i.e., for each connection separately) one-sample *t*-tests for the “baseline” effects of 3 Hz, 5 Hz, and 15 Hz, respectively. Furthermore, differences in effective connectivity patterns across the stimulation conditions were assessed using edgewise two-sample *t*-tests. To account for the number of tests performed (i.e., the number of connections), multiple comparison correction was based on the Benjamini-Hochberg procedure to control the false discovery rate (FDR).

#### Time-resolved network analyses

To inspect functional connectivity between cortical and thalamic regions of interest, we used the Louvain modularity algorithm from the Brain Connectivity Toolbox on the functional connectivity edge weights to estimate community structure ^83^. As in previous work ^52^, the stability of the gamma parameter was determined through a permutation process and defined as 1.3 for all analyses. We used the partition from the modularity algorithm to estimate the participation coefficient, *PC*, from un-thresholded, weighted, and signed connectivity matrices ^83^. The participation coefficient quantifies the extent to which a region connects across all modules (i.e., between-module strength) – PC is close to 1 if its connections are uniformly distributed among all the modules and 0 if all of its links are within its own module. The mean *PC* scores were then compared across subjects for the different stimulation types, after first correcting for the baseline associations observed during pre-stimulation ‘resting state’ epochs.

#### Brain State Displacement and the Energy Landscape

To quantify the change in BOLD activity following LC stimulation, we calculated the mean-squared displacement (MSD) of the BOLD signal over time. The MSD is a measure of the deviation in BOLD activity, *x*(*t*) = [*x*_1_ (*t*), *x*_2_ (*t*), …, *x*_*r*_ (*t*)] for *r* ROIs across time (*t*). The MSD is calculated as the mean of the squared change of each parcel’s activity over a time-delay, *dt*, given by

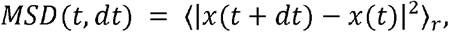

where ⟨ ⟩_*r*_ is the mean across *r* parcels. As in ^31^ we define the energy of observing a given brain-state change, *E*_*state*_, as the natural logarithm of its inverse probability, *P*_*state*_, thus,

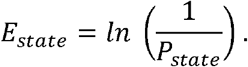

We estimated the probability distribution *P*_*state*_, from the sampled *MSD* (*t, dt*) within each of the experimental protocols using a Gaussian kernel density estimation 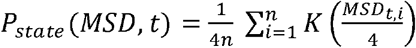, where 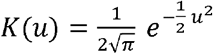 and we display the results for *dt* between 2 to 30 TR (2 to 30 s) and *MSD* between 0 to 10. The mean stimulation and post-stimulation energy landscapes were calculated across subjects for the different stimulation strengths, after first correcting for the individual subject baseline associations observed during pre-stimulation ‘resting state’ epochs.

## Notes

### Competing Interest Statement

The authors have declared no competing interest.

## References

1. Shine, J.M., et al. Human cognition involves the dynamic integration of neural activity and neuromodulatory systems. Nature Neuroscience 22, 289–296 (2019).

2. Shine, J. Neuromodulatory Influences on Integration and Segregation in the Brain. Trends in cognitive sciences 23 (2019).

3. Shine, J.M., et al. Computational models link cellular mechanisms of neuromodulation to large-scale neural dynamics. Nature Neuroscience 24, 765–776 (2021).

4. Aston-Jones, G. & Cohen, J.D. Adaptive gain and the role of the locus coeruleus-norepinephrine system in optimal performance. The Journal of comparative neurology 493 (2005).

5. Bouret, S. & Sara, S.J. Network reset: a simplified overarching theory of locus coeruleus noradrenaline function. Trends in neurosciences 28 (2005).

6. Yu, A.J. & Dayan, P. Uncertainty, neuromodulation, and attention. Neuron 46 (2005).

7. Berridge, C.W. & Waterhouse, B.D. The locus coeruleus-noradrenergic system: modulation of behavioral state and state-dependent cognitive processes. Brain research. Brain research reviews 42 (2003).

8. Poe, G.R., et al. Locus coeruleus: a new look at the blue spot. Nature Reviews Neuroscience 21, 644–659 (2020).

9. Totah, N., Logothetis, N. & Eschenko, O. Noradrenergic ensemble-based modulation of cognition over multiple timescales. Brain research 1709 (2019).

10. Bouret, S. & Sara, S.J. Reward expectation, orientation of attention and locus coeruleus-medial frontal cortex interplay during learning. European Journal of Neuroscience 20, 791–802 (2004).

11. Rajkowski, J., Majczynski, H., Clayton, E. & Aston-Jones, G. Activation of monkey locus coeruleus neurons varies with difficulty and performance in a target detection task. Journal of neurophysiology 92 (2004).

12. Vankov, A., Hervé-Minvielle, A. & Sara, S. Response to novelty and its rapid habituation in locus coeruleus neurons of the freely exploring rat. The European journal of neuroscience 7 (1995).

13. Takeuchi, T., et al. Locus coeruleus and dopaminergic consolidation of everyday memory. Nature 537 (2016).

14. Clayton, E.C. Phasic Activation of Monkey Locus Ceruleus Neurons by Simple Decisions in a Forced-Choice Task. Journal of Neuroscience 24, 9914–9920 (2004).

15. Aston-Jones, G. & Bloom, F. Nonrepinephrine-containing locus coeruleus neurons in behaving rats exhibit pronounced responses to non-noxious environmental stimuli. The Journal of Neuroscience 1, 887–900 (1981).

16. Mather, M., Clewett, D., Sakaki, M. & Harley, C. Norepinephrine ignites local hotspots of neuronal excitation: How arousal amplifies selectivity in perception and memory. The Behavioral and brain sciences 39 (2016).

17. Stellar, E. The physiology of motivation. 1954. Psychological review 101 (1994).

18. Sinnamon, H. Preoptic and hypothalamic neurons and the initiation of locomotion in the anesthetized rat. Progress in neurobiology 41 (1993).

19. Haberly, L. Parallel-distributed processing in olfactory cortex: new insights from morphological and physiological analysis of neuronal circuitry. Chemical senses 26 (2001).

20. Jordan, W. & Leaton, R. Startle habituation in rats after lesions in the brachium of the inferior colliculus. Physiology & behavior 28 (1982).

21. Avery, M. & Krichmar, J. Neuromodulatory Systems and Their Interactions: A Review of Models, Theories, and Experiments. Frontiers in neural circuits 11 (2017).

22. Lee, S.-H. & Dan, Y. Neuromodulation of Brain States. Neuron 76, 209–222 (2012).

23. Unsworth, N. & Robison, M.K. A locus coeruleus-norepinephrine account of individual differences in working memory capacity and attention control. Psychonomic Bulletin & Review 24, 1282–1311 (2017).

24. Berridge, C. & Abercrombie, E. Relationship between locus coeruleus discharge rates and rates of norepinephrine release within neocortex as assessed by in vivo microdialysis. Neuroscience 93 (1999).

25. Sara, J. Susan & Bouret, S. Orienting and Reorienting: The Locus Coeruleus Mediates Cognition through Arousal. Neuron 76, 130–141 (2012).

26. Carter, M.E., et al. Tuning arousal with optogenetic modulation of locus coeruleus neurons. Nature Neuroscience 13, 1526–1533 (2010).

27. Devilbiss, D. Consequences of tuning network function by tonic and phasic locus coeruleus output and stress: Regulating detection and discrimination of peripheral stimuli. Brain research 1709 (2019).

28. Hermans, E., et al. Stress-related noradrenergic activity prompts large-scale neural network reconfiguration. Science (New York, N.Y.) 334 (2011).

29. Ej, H., Mj, H., m, J. & G, F. Dynamic adaptation of large-scale brain networks in response to acute stressors. Trends in neurosciences 37 (2014).

30. Hasenkamp, W., Wilson-Mendenhall, C., Duncan, E. & Barsalou, L. Mind wandering and attention during focused meditation: a fine-grained temporal analysis of fluctuating cognitive states. NeuroImage 59 (2012).

31. Munn, B., Müller, E., Wainstein, G. & Shine, J. The ascending arousal system shapes neural dynamics to mediate awareness of cognitive states. Nature communications 12 (2021).

32. Zerbi, V., et al. Rapid Reconfiguration of the Functional Connectome after Chemogenetic Locus Coeruleus Activation. Neuron 103, 702-718.e705 (2019).

33. Oyarzabal, E., et al. Chemogenetic stimulation of tonic locus coeruleus activity strengthens the default mode network. Science advances 8 (2022).

34. Privitera, M., et al. A complete pupillometry toolbox for real-time monitoring of locus coeruleus activity in rodents. Nature Protocols 15, 2301–2320 (2020).

35. Bari, A., et al. Differential attentional control mechanisms by two distinct noradrenergic coeruleo-frontal cortical pathways. Proceedings of the National Academy of Sciences of the United States of America 117 (2020).

36. Rajkowski, J., Kubiak, P. & Aston-Jones, G. Locus coeruleus activity in monkey: phasic and tonic changes are associated with altered vigilance. Brain research bulletin 35 (1994).

37. Usher, M., Cohen, J., Servan-Schreiber, D., Rajkowski, J. & Aston-Jones, G. The role of locus coeruleus in the regulation of cognitive performance. Science (New York, N.Y.) 283 (1999).

38. McCall, J., et al. CRH Engagement of the Locus Coeruleus Noradrenergic System Mediates Stress-Induced Anxiety. Neuron 87 (2015).

39. Quinlan, M., et al. Locus Coeruleus Optogenetic Light Activation Induces Long-Term Potentiation of Perforant Path Population Spike Amplitude in Rat Dentate Gyrus. Frontiers in systems neuroscience 12 (2019).

40. Robertson, S.D., Plummer, N.W., de Marchena, J. & Jensen, P. Developmental origins of central norepinephrine neuron diversity. Nature Neuroscience 16, 1016–1023 (2013).

41. Feng, J., et al. A Genetically Encoded Fluorescent Sensor for Rapid and Specific In Vivo Detection of Norepinephrine. Neuron 102, 745-761.e748 (2019).

42. Ioanas, H.-I., Saab, B.J. & Rudin, M. Whole-brain opto-fMRI map of mouse VTA dopaminergic activation reflects structural projections with small but significant deviations. Translational Psychiatry 12, 1–10 (2022).

43. Aston-Jones, G., Rajkowski, J. & Cohen, J. Role of locus coeruleus in attention and behavioral flexibility. Biological psychiatry 46 (1999).

44. Cohen, R.A. Yerkes–Dodson Law. in Encyclopedia of Clinical Neuropsychology (ed. J.S. Kreutzer, J. DeLuca & B. Caplan) 2737–2738 (Springer New York, New York, NY, 2011).

45. Frässle, S., et al. Regression DCM for fMRI. NeuroImage 155 (2017).

46. Aston-Jones, G. & Cohen, J. An integrative theory of locus coeruleus-norepinephrine function: adaptive gain and optimal performance. Annual review of neuroscience 28 (2005).

47. McCormick, D., Nestvogel, D. & He, B. Neuromodulation of Brain State and Behavior. Annual review of neuroscience 43 (2020).

48. Shine, J. The thalamus integrates the macrosystems of the brain to facilitate complex, adaptive brain network dynamics. Progress in neurobiology 199 (2021).

49. Hwang, K., Bertolero, M., Liu, W. & D’Esposito, M. The Human Thalamus Is an Integrative Hub for Functional Brain Networks. The Journal of neuroscience : the official journal of the Society for Neuroscience 37 (2017).

50. Müller, E., et al. Core and matrix thalamic sub-populations relate to spatio-temporal cortical connectivity gradients. NeuroImage 222 (2020).

51. McCormick, D., McGinley, M. & Salkoff, D. Brain state dependent activity in the cortex and thalamus. Current opinion in neurobiology 31 (2015).

52. Shine, J., et al. The Dynamics of Functional Brain Networks: Integrated Network States during Cognitive Task Performance. Neuron 92 (2016).

53. Shine, J., Aburn, M., Breakspear, M. & Poldrack, R. The modulation of neural gain facilitates a transition between functional segregation and integration in the brain. eLife 7 (2018).

54. Fuxe, K., et al. The discovery of central monoamine neurons gave volume transmission to the wired brain. Progress in neurobiology 90 (2010).

55. Ghosh, A., et al. Locus Coeruleus Activation Patterns Differentially Modulate Odor Discrimination Learning and Odor Valence in Rats. Cerebral cortex communications 2 (2021).

56. Florin-Lechner, S., Druhan, J., Aston-Jones, G. & Valentino, R. Enhanced norepinephrine release in prefrontal cortex with burst stimulation of the locus coeruleus. Brain research 742 (1996).

57. McGinley, M., David, S. & McCormick, D. Cortical Membrane Potential Signature of Optimal States for Sensory Signal Detection. Neuron 87 (2015).

58. Aston-Jones, G., Rajkowski, J., Kubiak, P. & Alexinsky, T. Locus coeruleus neurons in monkey are selectively activated by attended cues in a vigilance task. The Journal of neuroscience : the official journal of the Society for Neuroscience 14 (1994).

59. Mccall, J.G., et al. Locus coeruleus to basolateral amygdala noradrenergic projections promote anxiety-like behavior. eLife 6 (2017).

60. Chandler, D., Waterhouse, B. & Gao, W. New perspectives on catecholaminergic regulation of executive circuits: evidence for independent modulation of prefrontal functions by midbrain dopaminergic and noradrenergic neurons. Frontiers in neural circuits 8 (2014).

61. Gompf, H.S., et al. Locus Ceruleus and Anterior Cingulate Cortex Sustain Wakefulness in a Novel Environment. Journal of Neuroscience 30, 14543–14551 (2010).

62. Arnsten, A. Through the looking glass: differential noradenergic modulation of prefrontal cortical function. Neural plasticity 7 (2000).

63. O’Rourke, M.F., Blaxall, H.S., Iversen, L.J. & Bylund, D.B. Characterization of [3H]RX821002 binding to alpha-2 adrenergic receptor subtypes. Journal of Pharmacology and Experimental Therapeutics 268, 1362–1367 (1994).

64. Arnsten, A., Cai, J. & Goldman-Rakic, P. The alpha-2 adrenergic agonist guanfacine improves memory in aged monkeys without sedative or hypotensive side effects: evidence for alpha-2 receptor subtypes. The Journal of neuroscience : the official journal of the Society for Neuroscience 8 (1988).

65. Mohell, N., Svartengren, J. & Cannon, B. Identification of [3H]prazosin binding sites in crude membranes and isolated cells of brown adipose tissue as alpha 1-adrenergic receptors. European journal of pharmacology 92 (1983).

66. Arnsten, A. & Goldman-Rakic, P. Noise stress impairs prefrontal cortical cognitive function in monkeys: evidence for a hyperdopaminergic mechanism. Archives of general psychiatry 55 (1998).

67. Arnsten, A., Mathew, R., Ubriani, R., Taylor, J. & Li, B. Alpha-1 noradrenergic receptor stimulation impairs prefrontal cortical cognitive function. Biological psychiatry 45 (1999).

68. Newman, L., Darling, J. & McGaughy, J. Atomoxetine reverses attentional deficits produced by noradrenergic deafferentation of medial prefrontal cortex. Psychopharmacology 200 (2008).

69. McGaughy, J., Ross, R. & Eichenbaum, H. Noradrenergic, but not cholinergic, deafferentation of prefrontal cortex impairs attentional set-shifting. Neuroscience 153 (2008).

70. Chandler, D., Lamperski, C. & Waterhouse, B. Identification and distribution of projections from monoaminergic and cholinergic nuclei to functionally differentiated subregions of prefrontal cortex. Brain research 1522 (2013).

71. Devilbiss, D.M. The Effects of Tonic Locus Ceruleus Output on Sensory-Evoked Responses of Ventral Posterior Medial Thalamic and Barrel Field Cortical Neurons in the Awake Rat. Journal of Neuroscience 24, 10773–10785 (2004).

72. Devilbiss, D., Page, M. & Waterhouse, B. Locus ceruleus regulates sensory encoding by neurons and networks in waking animals. The Journal of neuroscience : the official journal of the Society for Neuroscience 26 (2006).

73. Rodenkirch, C., Liu, Y., Schriver, B. & Wang, Q. Locus coeruleus activation enhances thalamic feature selectivity via norepinephrine regulation of intrathalamic circuit dynamics. Nature neuroscience 22 (2019).

74. Totah, N., Neves, R., Panzeri, S., Logothetis, N. & Eschenko, O. The Locus Coeruleus Is a Complex and Differentiated Neuromodulatory System. Neuron 99 (2018).

75. Uematsu, A., et al. Modular organization of the brainstem noradrenaline system coordinates opposing learning states. Nature Neuroscience 20, 1602–1611 (2017).

76. Chandler, D., Gao, W. & Waterhouse, B. Heterogeneous organization of the locus coeruleus projections to prefrontal and motor cortices. Proceedings of the National Academy of Sciences of the United States of America 111 (2014).

77. Parlato, R., Otto, C., Begus, Y., Stotz, S. & SchüTz, G.N. Specific ablation of the transcription factor CREB in sympathetic neurons surprisingly protects against developmentally regulated apoptosis. Development 134, 1663–1670 (2007).

78. Mathis, A., et al. DeepLabCut: markerless pose estimation of user-defined body parts with deep learning. Nat Neurosci 21, 1281–1289 (2018).

79. Nath, T., et al. Using DeepLabCut for 3D markerless pose estimation across species and behaviors. Nature protocols 14 (2019).

80. Frässle, S., et al. TAPAS: An Open-Source Software Package for Translational Neuromodeling and Computational Psychiatry. Frontiers in psychiatry 12 (2021).

81. Friston, K., Harrison, L. & Penny, W. Dynamic causal modelling. NeuroImage 19 (2003).

82. Frässle, S., et al. A generative model of whole-brain effective connectivity. NeuroImage 179 (2018).

83. Rubinov, M. & Sporns, O. Complex network measures of brain connectivity: uses and interpretations. NeuroImage 52 (2010).

