## Supplementary material for "Locus Coeruleus firing patterns selectively modulate brain activity and dynamics"

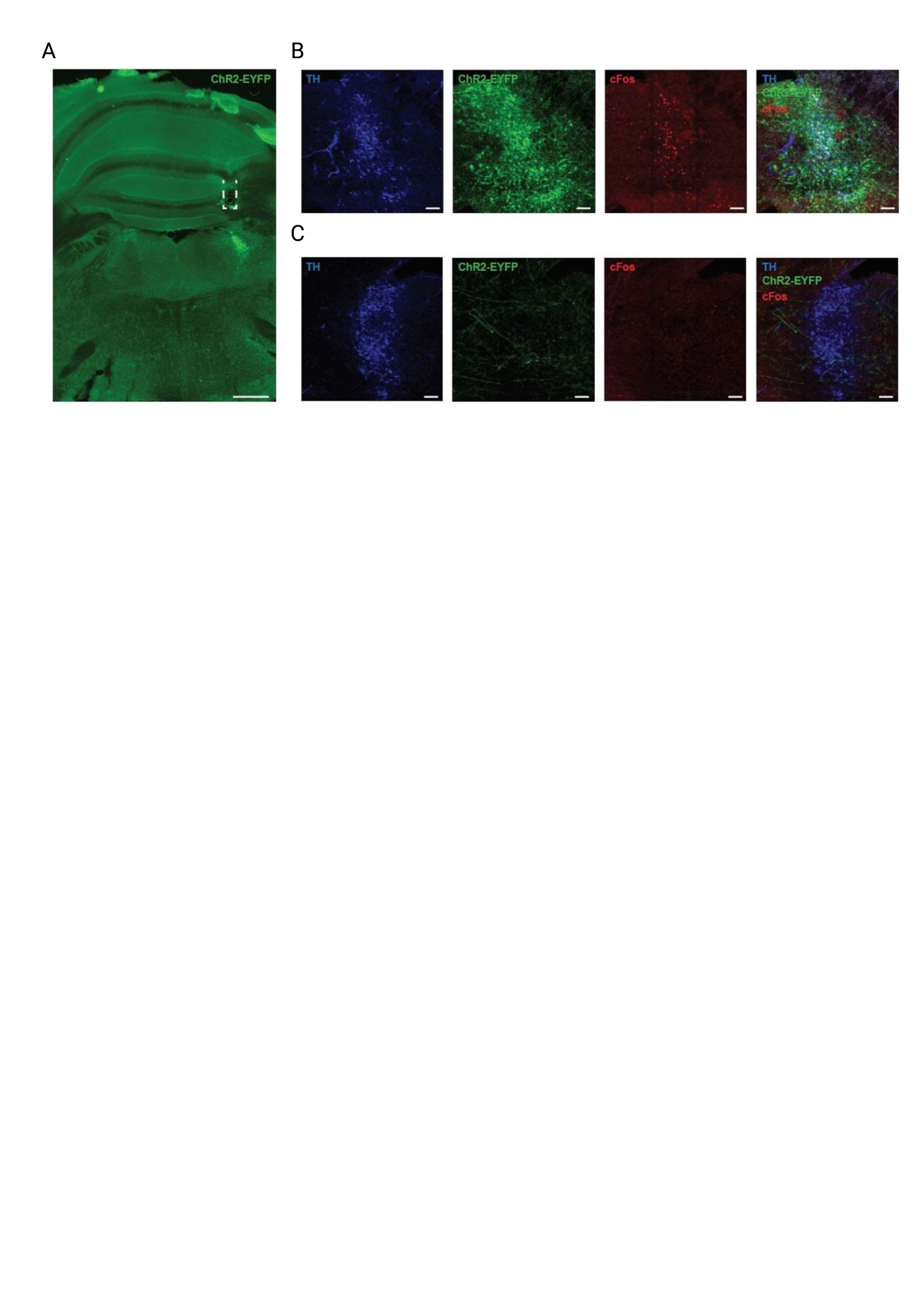

**Figure S1 Optogenetic targeting of LC noradrenergic neurons.** Related to Figure 3.1. **A** Representative immunohistochemical macro view of ChR2-EYFP targeting to the right (ipsilateral) LC. Dashes represent optical fibre placement. **B** Representative immunohistochemical view of ChR2-EYFP and cFos expression in TH-positive neurons of the right (stimulated) LC. **C** Representative immunohistochemical view of EYFP – positive axon fibres that project from the right (stimulated) to the left (unstimulated) LC. No cFos is detected in TH-positive neurons in the left LC. TH, tyrosine hydroxylase. Scale bars 500µm, 100µm.

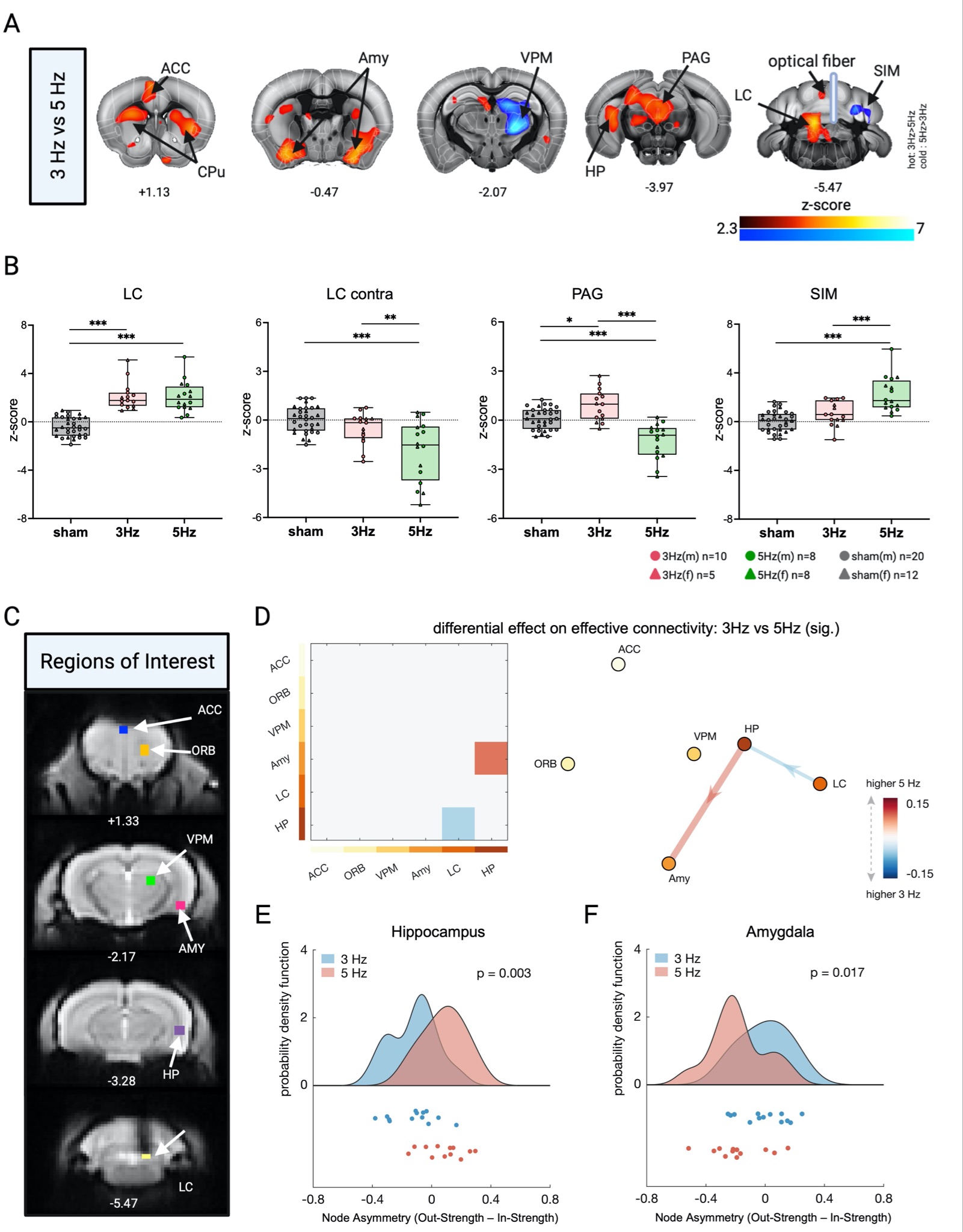

**Figure S2 Local effects of tonic LC stimulation.** Related to Figure 3. **A** Thresholded between-group statistical map of 3 Hz tonic vs 5 Hz tonic LC stimulation (cluster corrected; p<0.05). Colour code hot (red to bright yellow): 3Hz > 5Hz; colour code cold (blue to turquoise): 5Hz > 3Hz. **B** Mean Z-scores of male (circle) and female (triangle) mice in the ipsilateral LC significantly differed between stimulations (F(2,60)=48.790, p<0.0005). Mean Z-scores of male (circle) and female (triangle) mice in the contralateral LC significantly differed between stimulations (F(2,60)=13.670, p<0.0005). Mean Z-scores of male (circle) and female (triangle) mice in the ipsilateral PAG significantly differed between stimulations (F(2,60)=26.107, p<.0005). Mean Z-scores of male (circle) and female (triangle) mice in the ipsilateral SIM significantly differed between stimulations (F(2,60)=23.995, p<0.0005). Mean z-scores of sham datasets depicted in grey, mean z-scores of 3Hz tonic datasets depicted in red, mean z-scores of 5Hz tonic datasets depicted in green. **C** Selected regions of interest for rDCM analysis. Regions include the ACC (9 voxels, 200 mm3), ORB (12 voxels, 266 mm3), VPM (9 voxels, 200 mm3), amygdala (9 voxels, 200 mm3), HP (9 voxels, 200 mm3), and LC (8 voxels, 177 mm3). **D** Difference in the effective connectivity patterns between the tonic stimulation patterns (5Hz - 3Hz) in the six-region network. Only significant (p<0.05, FDR-corrected for multiple comparisons) differences are shown, both as an adjacency matrix (*left*) as well as a network graph (*right*). Warm colours indicate connections that were stronger during 5Hz as compared to 3Hz stimulation, and, vice versa, cool colours indicate connections that were stronger during 3Hz as compared to 5Hz stimulation. **E** Node asymmetry, quantified as the difference between out-strength (i.e., the sum of the strength of all outgoing connections) and in-strength (i.e., the sum of the strength of all ingoing connections), for the HP, as well as **F** for the Amy for the 3Hz (blue) and 5Hz (red) stimulation condition. HP showed significantly larger node asymmetry during the 5Hz as compared to 3Hz stimulation condition. Vice versa, Amy showed significantly larger node asymmetry during the 3Hz as compared to 5Hz stimulation condition. ACC, anterior cingulate; ORB, orbitofrontal cortex; CPu, caudate putamen; Amy, amygdala; HP, hippocampus PAG, periaqueductal gray; LC, locus coeruleus; SIM, simple lobule; VPM, ventral posteriomedial nucleus of thalamus; TH, thalamus; SSp, primary somatosensory cortex; SSs, supplemental somatosensory cortex. N(sham)=32; n(3Hz)=15; n(5Hz)=16.

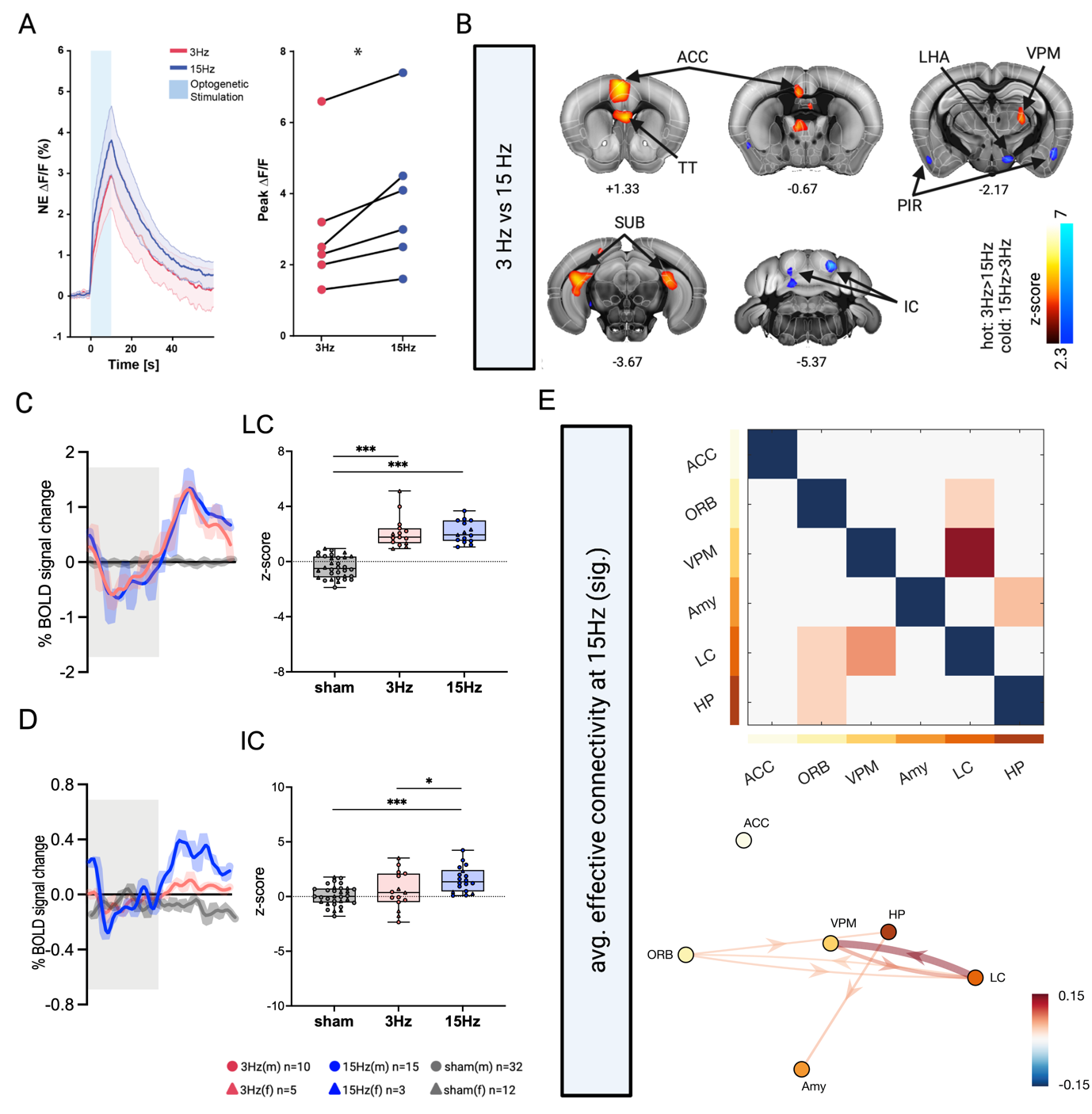

**Figure S3 Effects of 15 Hz burst vs 3 Hz tonic LC stimulation.** Related to Figure 4. **A** Left: ΔF/F traces of GRAB_NE1m_ photometry recordings in response to tonic 3Hz tonic and 15Hz phasic LC stimulation (mean±SEM). Right: 15Hz phasic LC stimulation triggers greater NA release compared to 3Hz tonic LC stimulation (mean±SD(3Hz) = 2.98 ± 1.88, mean±SD(15Hz) = 3.85 ± 2.03, paired t-test; t(5)=3.57, p=0.016). **B** Thresholded between-group statistical map of 3Hz tonic vs 15Hz burst LC stimulation (cluster corrected; p<0.05). Colour code hot (red to bright yellow): 3Hz > 15Hz; colour code cold (blue to turquoise): 15Hz > 3Hz. **C** Mean time-series extracted from targeted LC of sham, 3Hz tonic, and 15Hz burst datasets. Similar to 3Hz and 5Hz tonic LC stimulation, the BOLD response evoked by LC burst stimulation shows an initial dip followed by a delayed increase. This likely reflects not only the neural but also the neurovascular effects of NA release. Mean Z-scores of male (circle) and female (triangle) mice in the ipsilateral LC significantly differed between stimulations (F(2,62)=47.732, p<0.0005). **D** Mean time-series of inferior colliculus of sham, 3Hz tonic, and 15Hz burst datasets. Mean Z-scores of male (circle) and female (triangle) mice in the ipsilateral IC significantly differed between stimulations (F(2,62)=27.032, p<0.0005). Time-series of 3Hz tonic stimulation depicted in salmon, time-series of 15Hz tonic stimulation depicted in blue, time-series of sham depicted in grey. Grey bar represents 30s laser stimulation block. **E** Average connectivity pattern during the 15Hz bust stimulation condition. Only significant (p<0.05, FDR-corrected for multiple comparisons) connections are shown, both as an adjacency matrix (*top*) as well as a network graph (*bottom*). LC, locus coeruleus; VPM, ventral posteriomedial nucleus of thalamus; TT, taenia tecta; ACC, anterior cingulate; ORB, orbitofrontal area; Amy, amygdala; HP, hippocampus; SUB, subiculum; LHA, lateral hypothalamic area; PIR, piriform area; IC, inferior colliculus. N(sham)=32; n(3Hz)=15; n(15Hz)=18. *p < 0.05; ***p < 0.001

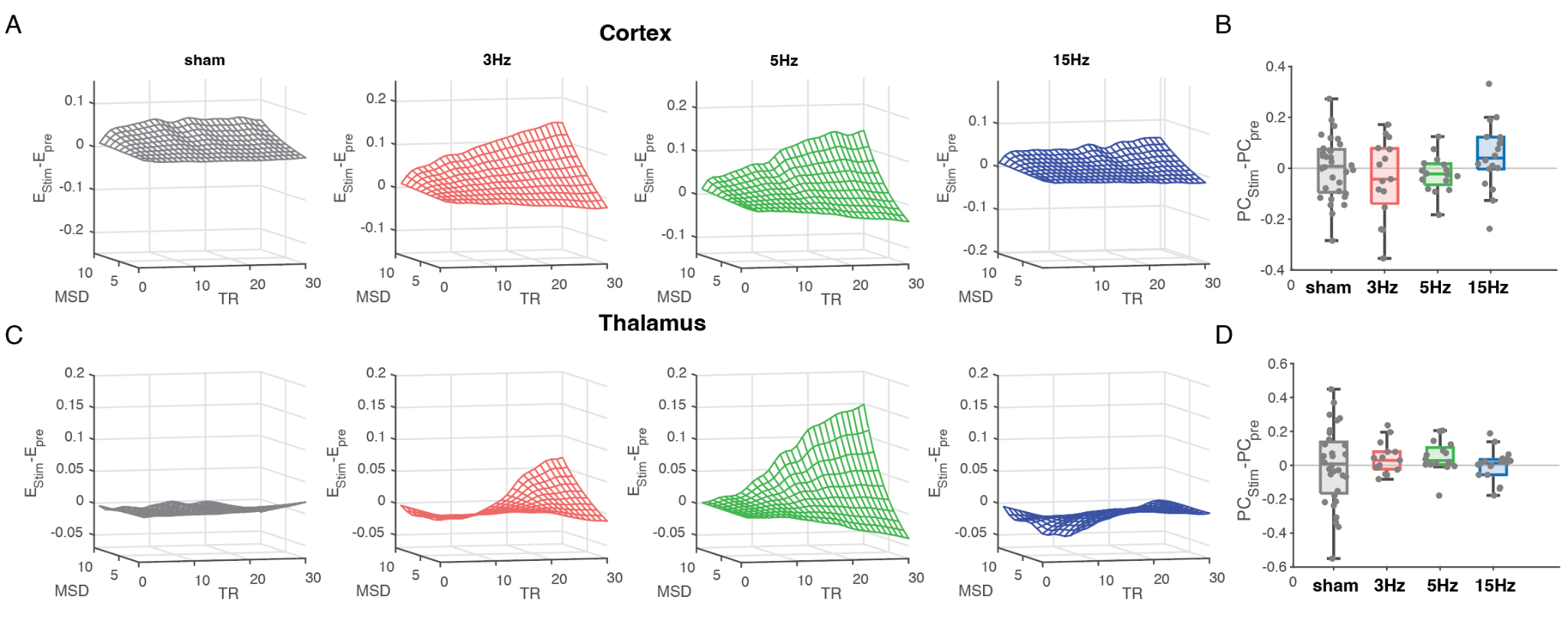

**Figure S4 Post stimulation energy landscape topography and network topology differ between tonic and phasic stimulation. A** Following stimulation cortical energy landscapes demonstrate a rebound exhaustion increasing the energy of large BOLD changes (3Hz (red) & 5Hz (green) vs sham both p<0.05, paired t-test, Bonferroni corrected) whereas phasic stimulation is indistinguishable from prestimulation regime (~0). **B** Network dynamics following tonic stimulation also show a rebound decrease in PC (i.e., increase in network segregation; p<0.05, SEM) whereas 15Hz burst stimulation leads to a prolonged cortical integration (p<0.05, SEM). **C** Thalamic energy landscapes 5Hz display a statistically significant difference between sham with a rebound of increased energy for large BOLD changes relative to prestimulation baseline (p<0.05, paired t-test). **D** Thalamic network changes demonstrate prolonged integration following tonic but not phasic stimulation (p<0.05, SEM).
